# Harnessing CRISPR interference to re-sensitize laboratory strains and clinical isolates to last resort antibiotics

**DOI:** 10.1101/2024.07.30.604783

**Authors:** Angelica Frusteri Chiacchiera, Michela Casanova, Massimo Bellato, Aurora Piazza, Roberta Migliavacca, Gregory Batt, Paolo Magni, Lorenzo Pasotti

**Affiliations:** Department of Electrical, Computer and Biomedical Engineering, University of Pavia, Via Ferrata 5, Pavia, Italy; Centre for Health Technologies, University of Pavia, Via Ferrata 5, Pavia, Italy; Institut Pasteur, Inria, Université Paris Cité, 28 rue du Docteur Roux, Paris, France; Department of Information Engineering, University of Padua, Via Gradenigo 6b, 35131 Padua, Italy; Department of Clinical-Surgical, Diagnostic and Pediatric Sciences, University of Pavia, Viale Brambilla 74, Pavia, Italy; Fondazione IRCCS Policlinico San Matteo, Pavia, Italy

**Keywords:** Antimicrobial resistance, antibiotic re-sensitization, CRISPR array, *Escherichia coli* clinical isolates, *bla*_NDM_-type, *bla*_ctx-M_-type, *mcr-1*

## Abstract

The global race against antimicrobial resistance requires novel antimicrobials that are not only effective in killing specific bacteria, but also minimize the emergence of new resistances. Recently, CRISPR/Cas-based antimicrobials were proposed to address killing specificity with encouraging results. However, the emergence of target sequence mutations triggered by Cas-cleavage was identified as an escape strategy, posing the risk of generating new antibiotic-resistance gene (ARG) variants. Here, we evaluated an antibiotic re-sensitization strategy based on CRISPR interference (CRISPRi), which inhibits gene expression without damaging target DNA. The resistance to four antibiotics, including last resort drugs, was significantly reduced by individual and multi-gene targeting of ARGs in low- to high-copy numbers in recombinant *E. coli*. Escaper analysis confirmed the absence of mutations in target sequence, corroborating the harmless role of CRISPRi in the selection of new resistances. *E. coli* clinical isolates carrying ARGs of severe clinical concern were then used to test the robustness of CRISPRi under different growth conditions. Meropenem, colistin and cefotaxime susceptibility was successfully increased in terms of MIC (up to >4-fold) and growth delay (up to 11-hours) in a medium-dependent fashion. To our knowledge, this is the first demonstration of CRISPRi-mediated re-sensitization to last-resort drugs in clinical isolates. This study laid the foundations for further leveraging CRISPRi as antimicrobial agent or research tool to selectively repress ARGs and investigate resistance mechanisms.

## Introduction

The rapid spread of multidrug resistant pathogens has become a global health concern, exacerbated by the lack of innovation displayed by the clinical and pre-clinical pipelines for developing new antibiotics^1,2^. Alternatives to traditional antibiotic treatments are thus urgently needed to eradicate antimicrobial resistance (AMR)-associated infections and minimize the evolution of antibiotic-escape mechanisms. In recent years, CRISPR-based approaches have been proposed to develop both therapeutic agents acting as alternative or adjuvant to antibiotics^3–5^, and molecular genetic tools for research purposes^6–13^. For instance, CRISPR antimicrobials have been designed by redirecting the cleavage of Cas nucleases towards chromosome-encoded or plasmid-borne antibiotic resistance genes (ARGs), resulting in cell death or antibiotic re-sensitization, respectively^14–17^. Regarding CRISPR delivery into target cells, it has been addressed with different vehicles^12,14–20^, including conjugative plasmids that can be optimized to reach almost 100% conjugation efficiency in the mouse gut^21^.

These studies have also unearthed the emergence of escape mutants. In particular, as a consequence of the SOS response triggered by Cas9-mediated double strand breaks (DSB), mutations can occur in the CRISPR circuitry or in the target gene^19,20,22–24^. Indeed, it has been reported that escapers can occur at a rate higher than the one considered as acceptable by the National Institute of Health^25^, hindering the application of CRISPR-based antimicrobials in clinical settings. In the field of antimicrobial resistance, CRISPR-escape mutants that have evolved target gene mutations are of specific concern since the new generated sequence became immune to CRISPR targeting and may still encode a functional protein, thus representing a new resistance variant. To counteract this risk, the CRISPR *interference* (CRISPRi) technology could represent an attractive solution as it relies on the ability of dead Cas9 (dCas9) to inhibit the expression of a desired gene without damaging the target DNA^26,27^. Re-sensitized cells are eventually killed upon antibiotic administration. This approach aims to preserve the target gene sequence, while other strategies can be exploited to counteract mutations that inactivate the CRISPR circuit, such as the design of CRISPR arrays with multiple guides for the same gene to increase on-target efficiency.

To date, the CRISPRi-based approach has been evaluated against ARGs located on the chromosome or low copy plasmids. Laboratory strains of *Escherichia coli* or *Mycobacterium smegmatis* have been re-sensitized to ampicillin, trimethoprim, sulfamethoxazole, or rifampicin, respectively^20,28,29^. To our knowledge, only one study addresses the CRISPRi-based antibiotic re-sensitization of a clinical isolate, a methicillin-resistant *Staphylococcus aureus*^30^. Beyond the antimicrobial function, CRISPRi-based tools have been exploited to perform gene functional screenings for validating drug targets, dissecting AMR mechanisms by systematically perturbing the expression of single genes and gene combinations, also gaining insight into the consequences of partial gene inhibition^8–11,31^. Although the mentioned studies highlight the potential of CRISPRi in both the design of antimicrobials and systems biology tools, a number of challenges still need to be addressed. The repression capability of the dCas9:sgRNA complex must be demonstrated even towards target genes present on high copy plasmids^32^ (I). Since the expression of a given ARG can be driven by different promoters in different strains, it is necessary to demonstrate the efficiency of CRISPRi systems targeting CDS to support their application in real case studies (II). Although dCas9-mediated silencing is not expected to induce mutations in target genes, a characterization of the escaper cells still needs to be carried out in order to investigate the evolution of escape mutations (III). Multi-targeting of sequences within the same or different ARGs is a convenient feature, e.g., to selectively repress multiple resistances, and needs to be investigated. Also, demonstrative work assessing the efficiency of CRISPRi platforms in pathogens carrying clinically relevant ARGs is desirable (IV). Finally, the robustness of the proposed solution needs to be assessed in different environments in order to quantify the impact of growth conditions on bacterial antibiotic response and CRISPRi-repression capability, and to evaluate its potential application for clinically-relevant studies (V). In this work, we have designed and characterized a CRISPRi-based platform circuitry to repress different ARGs and quantify the resulting effect on antibiotic re-sensitization. First, repression performance was evaluated and further optimized in recombinant *E. coli* strains expressing four ARGs (*bla*_TEM-116_, *tetA, bla*_NDM-1_, *mcr-1*), by testing their single and multiple targeting. These ARGs were selected as case studies due to their different gene dosages (low- to high-copy plasmids), resistance mechanisms, and therapeutic relevance in clinical settings. Analysis of escaper cells was carried out to study the bacterial response upon antibiotic re-exposure and to highlight the genetic mutations responsible for the rescue of surviving cells. Finally, to test the potential of our circuitry in clinically relevant case studies, we constructed a CRISPRi-based trans-conjugative platform to carry out a full delivery and re-sensitization workflow from engineered donors to four pathogenic *E. coli* strains: *bla*_NDM-5_, *mcr-1, bla*_ctx-M-14_ and *bla*_*c*tx-M-15_ were individually repressed and their growth inhibition effects characterized in three media providing different growth conditions.

## Results

### Repression of single and multiple ARGs in recombinant bacteria

The CRISPRi-mediated antibiotic re-sensitization was first assessed in recombinant *E. coli* strains. This preliminary investigation was conducted on ad-hoc constructed case studies to test re-sensitization of strains with ARG in high-copy plasmid as well as the simultaneous inhibition of two ARGs. Both studies have not been yet evaluated in the literature of CRISPRi antimicrobials. *tetA* or *bla*_*TEM-116*_ were used as target ARGs. *tetA* is located on a very low copy F’ episome (vLC, ∼1.5 copies) and encodes for an efflux pump conferring resistance to tetracycline (TC); *bla*_*TEM-116*_ is located on a high-copy vector (HC, ∼70 copies) and encodes a broad-spectrum β-lactamase which provides resistance to many β-lactam antibiotics, including ampicillin (AMP). Such targets were selected due to their different resistance mechanisms and to evaluate CRISPRi efficiency against plasmids with largely different copy numbers. A set of three test strains was investigated: *sensitive* strain, without ARG-carrying plasmids; *resistant* strain, with the target ARG(s); *specific CRISPRi transformant* (henceforth *sCRISPRi*), bearing the ARG-carrying plasmid(s) and a CRISPRi platform targeting one or two ARGs (Figure S1a). This set of strains was characterized via microplate and agar plate assays to quantify antibiotic re-sensitization in terms of minimum inhibitory concentration (MIC), growth delays (Δt) and inhibitory concentration killing at least 99% of the population (IC_99_) (Figure S1b).

### Tetracycline re-sensitization in microplate assays by single ARG targeting

A two-plasmid CRISPRi platform targeting single ARGs has been adopted (Figure 1a, Figure S1a). The platform relied on a low-copy plasmid (LC, ∼5 copies)^33^, carrying the gRNA module under the control of an isopropyl-β-D-1-thiogalactopyranoside (IPTG)-inducible P_LlacO1_ promoter, and a medium-copy plasmid (MC, ∼15 copies)^33,34^, carrying the dCas9 module under the P_J23116_ weak constitutive promoter, previously evaluated as a high-repression efficiency and low-burden dCas9 expression cassette^35^ (Figure 1a). A gRNA module (gtetA) was designed to block *tetA* transcription by targeting its CDS. Representative growth profiles at different TC concentrations are reported in Figure 1b for *resistant* and *sCRISPRi* strain and Figure S2a for *sensitive* strain. We observed that the MIC of the *resistant* strain (MIC>100 µg/ml) was more than 10-fold higher than that of the *sensitive* strain (MIC=10 µg/ml); nonetheless, the growth of resistant cells started to be compromised for TC>10 µg/ml, resulting in a delayed growth profile. On the other hand, *tetA* inhibition in the *sCRISPRi* strain contributed to a substantial increase in the time needed for population recovery in presence of sub-inhibitory TC concentrations, leading to a complete TC re-sensitization with a MIC of 100 µg/ml (Figure 1b). The delayed growth of the *sCRISPRi* strain was interpreted as a response to *tetA* inhibition by dCas9:gtetA, drug sequestration (TetR could still reduce the concentration of free intracellular TC^36^) and possible loss of function mutations occurring in the CRISPRi circuitry. The delays in *sCRISPRi* strain recovery increased as a function of TC concentration, with significantly larger delay values than the *resistant* strain (p<0.05, t-test) (Figure 1b). The *sCRISPRi* strain tested in the absence of IPTG indicated that the leakiness of P_LlacO1_ activity was sufficient to restore TC susceptibility, as a low amount of the dCas9:gtetA repressor complex was already effective in inhibiting *tetA* expressed from a vLC plasmid with no relevant difference from the on-state (Figure S3a,c).

**Figure 1.**
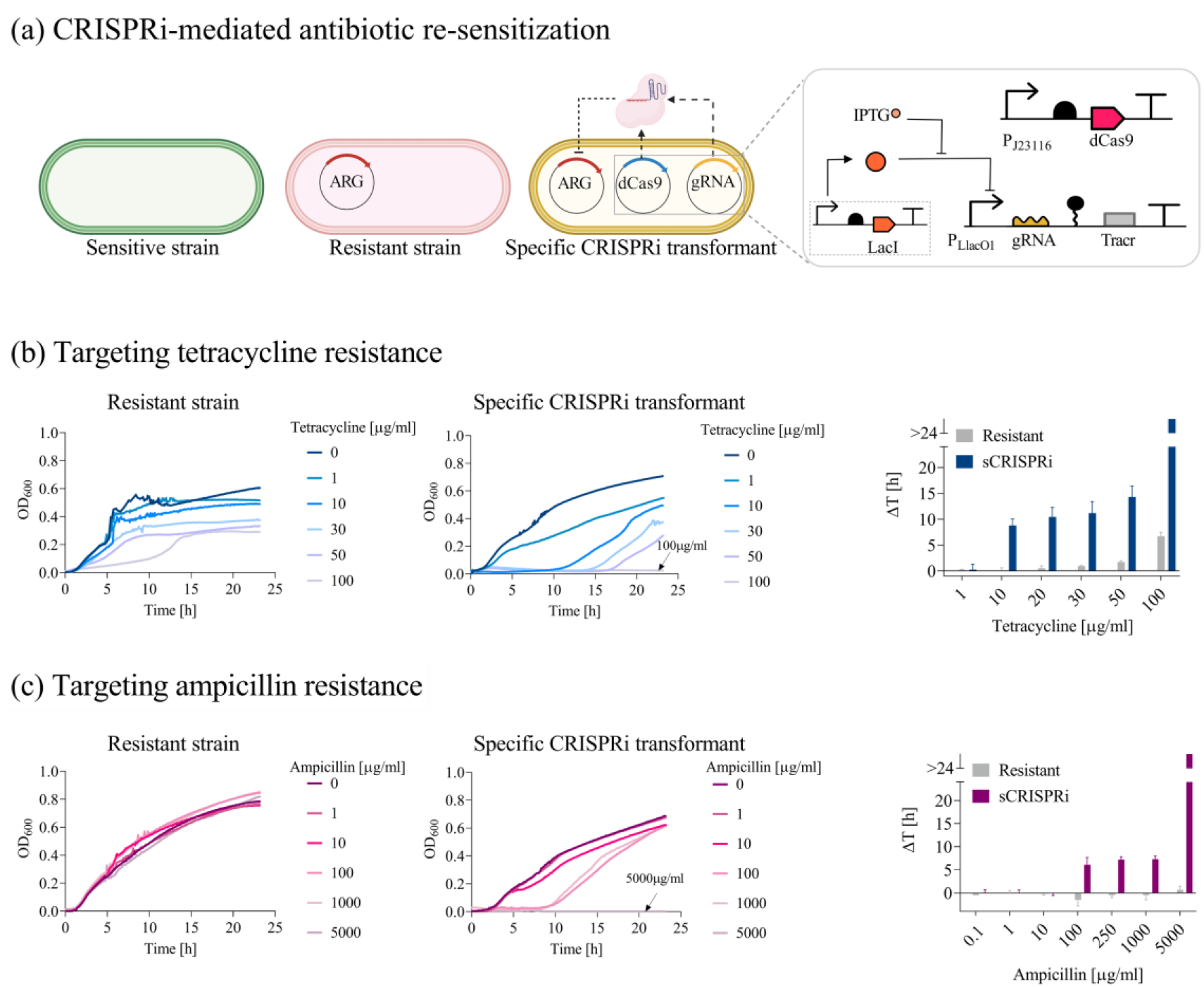
Re-sensitization of laboratory strains to tetracycline and ampicillin by single ARG targeting. **(a)** Description of the test strains. The inhibition of *tetA* or *bla*_TEM-116_ (red) is investigated with a two-plasmid CRISPRi platform, including a constitutive dCas9 module (blue) in a medium-copy plasmid and an IPTG-inducible sgRNA module (yellow) in a low-copy plasmid, transcribing gTetA, gbla_promoter_ or gbla_CDS_. Platform components are detailed in the inset according to the Synthetic Biology Open Language (SBOL) notation. **(b-c)** Growth profiles (subpanels on the left) and growth delays (subpanels on the right) of the *resistant* and *sCRISPRi* strains from microplate assays to investigate tetracycline (b) and ampicillin (c) re-sensitization. *sCRISPRi* strains are engineered with a CRISPRi platform, including constitutive dCas9 and IPTG-inducible gtetA or gbla_promoter_ targeting *tetA* or *bla*_TEM-116_, respectively. The overlapped curves in which a complete growth inhibition was achieved are indicated with an arrow, with the list of the corresponding antibiotic concentrations. Representative curves are shown from a set of at least three independent experiments. Growth delays (Δt) of treated strains are relative to the growth profile without antibiotics for the *resistant* and *sCRISPRi* strains. Bars represent means and standard deviations (N=3). The bars over the interrupted axis indicate the antibiotic concentration (MIC) for which OD_600_ was lower than 0.1. Data come from experiments in LB media in the presence of IPTG. TOP10F’ and T-cr_sgT_const_ were used as *resistant* and *sCRISPRi* strains, respectively, to investigate tetracycline re-sensitization. A-res and A-cr_sgA_const_ were used as *resistant* and *sCRISPRi* strains, respectively, to investigate ampicillin re-sensitization.

### Ampicillin re-sensitization in microplate assays by single ARG targeting

The same CRISPRi platform described in Figure 1a was used to test *bla*_TEM-116_ inhibition. Two gRNAs (gbla_promoter_ and gbla_CDS_) were designed to prevent or block *bla*_TEM-116_ transcription by targeting its promoter or CDS, respectively (Figure 1c, Figure S4). We observed that the *resistant* strain was able to withstand even the highest AMP concentration tested, without relevant growth defects, resulting in a MIC>5000 µg/ml (Figure 1c), which was much higher than that of the *sensitive* strain (MIC=10 µg/ml) (Figure S2b). In the *sCRISPRi* strain, *bla*_TEM-116_ inhibition by a gRNA targeting *bla* promoter led to a MIC of 5000 µg/ml and a growth delay of 6 to 7 h for AMP between 10 and 1000 µg/ml (Figure 1c). Targeting the *bla* CDS instead of its promoter had a weaker effect, since a complete growth inhibition was not observed for the antibiotic concentrations tested. A higher growth delay than the resistant control was detected only for AMP>250 µg/ml (p<0.05, t-test), reaching a 7-h delay only for AMP=5000 µg/ml (Figure S4). These data confirm that the location of the target site can strongly affect repression efficiency, as previously described^26^. As expected, growth profiles in the absence of IPTG showed that in the context of ARG in HC plasmid the repression performance was lower than in a vLC plasmid, tested above, as the leakage of P_LlacO1_ led to a slightly delayed growth (2.5 h) only for the highest AMP concentration (p<0.05, t-test) (Figure S4). Here, different mechanisms can contribute to the growth delays, including *bla*_TEM-116_ repression, occurrence of mutations inactivating CRISPRi circuitry or emergence of collective antibiotic tolerance^37^.

Another set of TC- and AMP-resistant control strains, bearing a full CRISPRi plasmid set with non-specific sgRNA (*nsCRISPRi* with gbla_CDS_ and gtetA, respectively), was evaluated, yielding similar delay patterns to the *resistant* strains described above (Figure S5). This demonstrates the specificity of our system and also that the two CRISPRi plasmids used in this section did not cause relevant burden to the engineered cells. Finally, escaper cells recovered from the highest AMP and TC concentration were used to perform a second round of antibiotic treatment. We observed comparable growth profiles with the *resistant* strain (Figure S6), suggesting that a population of mutants with increased MIC was selected over the 24 h time course.

### Tetracycline and ampicillin re-sensitization in agar plate assays by single ARG targeting

To test whether the growth in liquid or solid media could affect the rescue of escaper cells, we performed agar plate assays (Figure S1, S7, and Table S4). We expected different results from liquid cultures in which an enrichment of escapers can occur due to the activity of enzymes degrading the antibiotic in the media (i.e., β-lactamases released by lysed cells allow population recovery via a phenomenon known as collective antibiotic tolerance^37^). This response is likely to be mitigated for isolated colonies on agar plates. Data of TC and AMP agar plate assays were qualitatively consistent with the data in microplate assays (Figure S7 and Table S4): *sCRISPRi* strains (*sCRISPRi-sgRNA* in the figure) exhibited lower IC_99_ values (TC = 5 µg/ml, AMP = 50 µg/ml) than *resistant* strains (TC = 50 µg/ml, AMP > 1000 µg/ml), resulting in a 10-fold and >20-fold increase of antibiotic susceptibility for TC and AMP, respectively. Consistent with microplate assays, the gbla_CDS_-strain exhibited a higher IC_99_ (150 µg/ml) than the gbla_promoter_ strain, still representing a relevant improvement in growth inhibition (>6-fold susceptibility compared with *resistant* strain). These data confirm that our CRISPRi platform can effectively inhibit single ARGs in low but also HC plasmids, leading to increased sensitivity to the respective antibiotics in recombinant *E. coli*.

### Multi-targeting of ARGs using a constitutive dCas9

The platform described above was extended to address the simultaneous targeting of *tetA* and *bla*_*TEM-116*_. The individual sgRNA cassette was replaced with two guide RNA architectures: double sgRNA cassette or CRISPRi array^38-40^. Both of them enable the simultaneous transcription of gtetA and gbla_promoter_ under the control of the same regulatory elements (IPTG-inducible P_LlacO1_ promoter and synthetic terminator) (Figure 2a). The transcription of dCas9 protein was kept under the control of the P_J23116_ weak constitutive promoter, to eventually investigate the effect of dCas9 sharing between different guides on their multi-targeting capability (Figure 2a). The resulting *sCRISPRi* strains harbored a two-plasmid CRISPRi platform and two different ARG-carrying plasmids (Figure S1a, Table S1). The described platform showed a lower performance than the individual gRNA cassette tested above, as MICs could not be determined in liquid assays (Figure 2b,d, and Figure S8). The decrease in repression efficiency was interpreted as the result of gRNAs competition for the shared pool of dCas9 protein, which ultimately affects the number of repressor complexes that can bind and inhibit target genes^41,42^.

**Figure 2.**
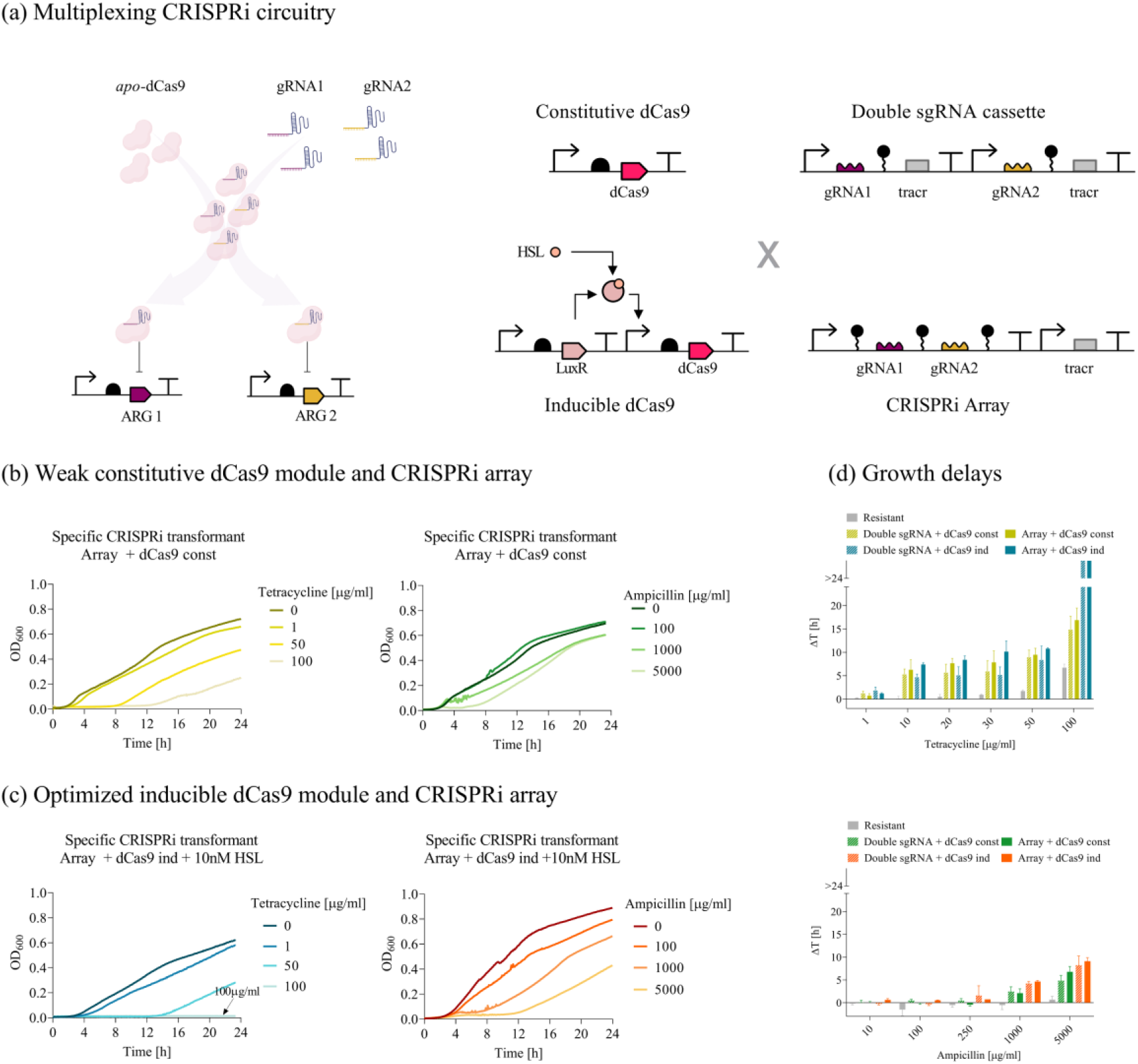
Re-sensitization of laboratory strains to tetracycline and ampicillin by multiple ARG targeting. **(a)** Illustration of multiplexed targeting of tetracycline and ampicillin resistance genes by two guides repressing *tetA* and *bla*_TEM-116_. Description of all the synthetic circuit combinations tested with two guide RNAs: weak constitutive or N-3-oxohexanoyl-L-homoserine lactone (HSL)-inducible dCas9, and IPTG-inducible CRISPRi array or sgRNA tandem cassettes (double sgRNA cassette), yielding four different circuit designs. **(b-c)** Growth profiles of *sCRISPRi* strains from microplate assays to investigate tetracycline (left subpanel) and ampicillin (right subpanel) re-sensitization. *sCRISPRi* strains are engineered with a two-plasmid CRISPRi platform, including constitutive dCas9 and IPTG-inducible CRISPRi array transcribing gtetA and gbla_promoter_ simultaneously (panel b). *sCRISPRi* strains are engineered with a one-plasmid CRISPRi platform, including HSL-inducible dCas9 (HSL concentration was set to 10 nM) and IPTG-inducible CRISPRi array transcribing gtetA and gbla_promoter_ simultaneously (panel c). Representative curves are shown in (b,c) from a set of at least three independent experiments. **(d)** Growth delays (Δt) of treated strains relative to the growth profile without antibiotics for the *resistant* strain and the four *sCRISPRi* strains with two-guide architectures. Bars represent means and standard deviations (N=3). The bars over the interrupted axis indicate the antibiotic concentration (MIC) for which OD_600_ was lower than 0.1. All the data come from experiments in LB media in the presence of IPTG and HSL. AT-cr_sgAT_const_ and AT-cr_arAT_const_ were used as *sCRISPRi* strains with constitutive dCas9 and inducible double sgRNA cassette or CRISPRi array, respectively. AT-cr_sgAT_ and AT-cr_arAT_ were used as *sCRISPRi* strains with inducible dCas9 and inducible double sgRNA cassette or CRISPRi array, respectively.

### Multi-targeting of ARGs using an inducible dCas9

To overcome the competition between gRNAs, an N-3-oxohexanoyl-L-homoserine lactone (HSL)-inducible dCas9 module was placed in the same plasmid already hosting the double sgRNA cassette or CRISPRi array, leading to the transition to a one-plasmid CRISPRi platform (Figure 2a, Figure S1a). TC and AMP growth inhibition assays were performed with a range of HSL concentrations to find an optimal dCas9 expression level, which was determined based on the analysis of growth delays (Figure S9). Regarding TC-treatment, a full susceptibility was restored in the HSL-inducible dCas9 strain at the same MIC as the single sgRNA cassette (MIC=100 µg/ml). Regarding AMP treatment, a major improvement over the previous platform is evidenced by the significantly increased delay in growth recovery at AMP concentrations higher than 250 µg/ml (p<0.05, t-test) (Figure 2c,d). Data from agar plate assays showed a consistent pattern (Figure S7): the one-plasmid inducible circuit displayed the same re-sensitization efficiency to TC as the single sgRNA cassette (IC_99_ =5 µg/ml); for AMP, a 4-fold increase of IC_99_ (200 µg/ml) was observed compared to that of the *sCRISPRi* strain bearing a single sgRNA cassette (50 µg/ml), confirming the impact of guide RNA competition on the silencing performances of a high copy number target gene. Overall, data showed that dCas9 tuning contributed to increase both TC- and AMP-sensitivity, although it was not sufficient to fully counteract gRNAs competition for HC-targets. Finally, the two gRNAs architectures were compared based on the analysis of growth delays at the fixed HSL concentration. Due to its better performance, the inducible dCas9 with CRISPRi array design was selected for subsequent analyses (Figure S9). This double targeting system also enabled to increase the recovery time of laboratory strains bearing both *bla*_TEM-116_ and *tetA* in a combined treatment with AMP and TET (see Figure S10).

## Sensitization of engineered bacteria to last-resort antibiotics

We reprogrammed the one-plasmid system with CRISPRi array described in Figure 2a to inhibit the expression of two clinically relevant ARGs, *bla*_NDM_ and *mcr*, conferring resistance to meropenem (MER) and colistin (COL), respectively. Both antibiotics are classified as last-resort drugs, representing an example of the few remaining therapeutic options available to treat severe AMR-associated infections. A set of four CRISPRi arrays was designed to target conserved positions among *bla*_NDM_ and *mcr* variants (Figure 3a and Figure S11). In particular, the inhibition of *bla*_NDM_ was characterized with two gRNAs (gNDM1 and gNDM2) included as single- (arM1, arM2) or double-spacers (arM1M2) in the CRISPRi array. The comparison between individual and combined spacers against *bla*_NDM_ was investigated to determine whether increasing the number of target sequences could enhance repression efficiency and/or limit escape mutations. The efficiency of a fourth array carrying a non-targeting gRNA (arM1C1) was also characterized for *bla*_NDM_ inhibition. The same array was used to test the inhibition of *mcr* with one gRNA (gMCR) (Figure 3a). MC vectors were transformed in *resistant* and *sCRISPRi* strains to express the ARGs (pGDP1 NDM-1 or pGDP2 MCR-1, bearing an *E. coli* codon-optimized *bla*_NDM-1_ driven by the P_bla_ promoter, and an *mcr-1* gene driven by the P_lac_ promoter)^43^ (Figure 3a).

**Figure 3.**
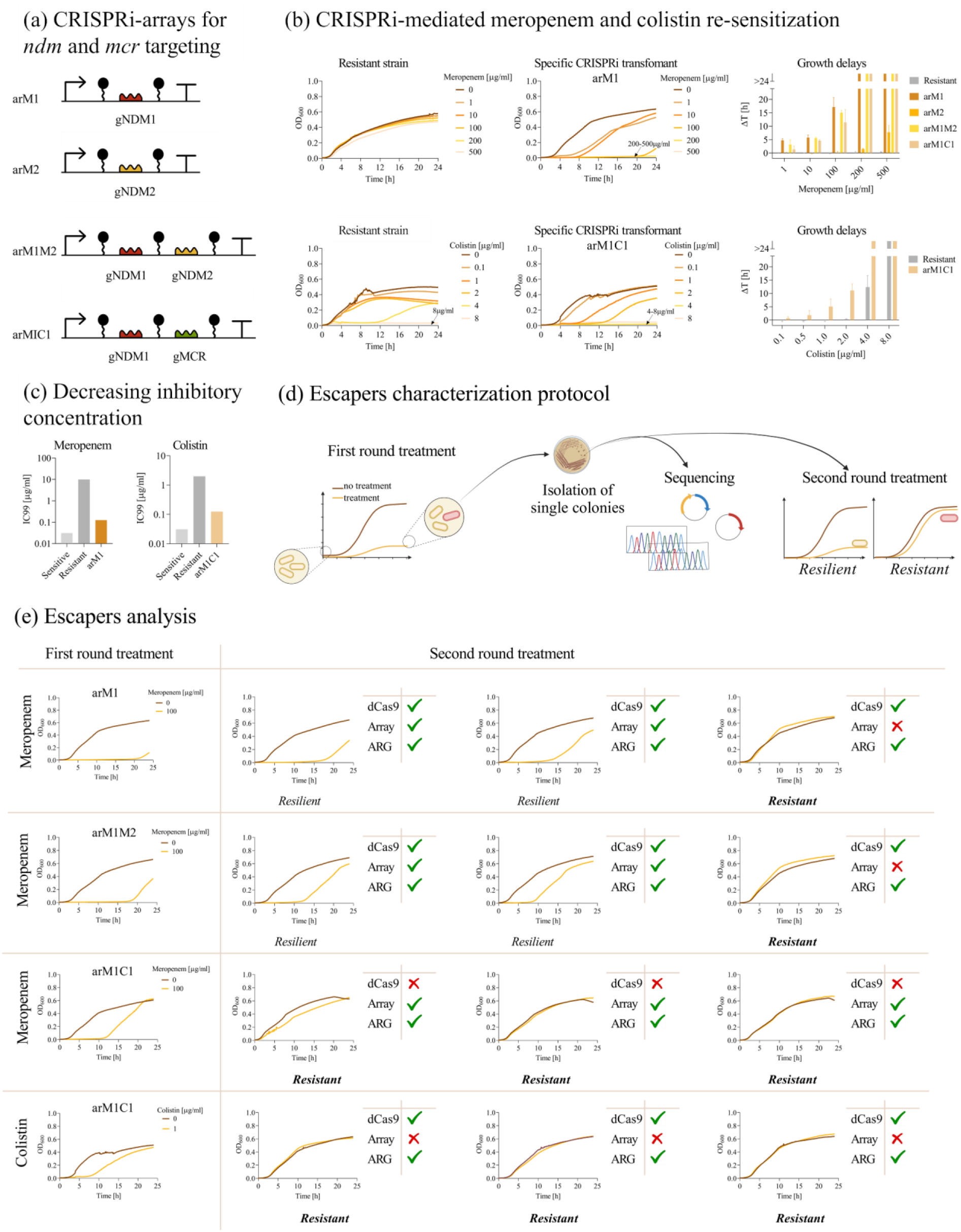
Re-sensitization of laboratory strains to last-resort antibiotics by single and multiple ARG targeting. **(a)** Description of CRISPRi arrays transcribing single or double spacers for *bla*_NDM-1_ and/or *mcr-1* targeting, with names provided on the left. **(b)** Growth profiles of *resistant* and *sCRISPRi* strains from microplate assays to investigate meropenem (top) and colistin (bottom) re-sensitization. *sCRISPRi* strains are engineered with a one-plasmid CRISPRi platform, including HSL-inducible dCas9 and IPTG-inducible CRISPRi arrays targeting *bla*_NDM-1_ (arM1) or *mcr-1* (arM1C1) gene. The overlapped curves in which a complete growth inhibition was achieved are indicated with an arrow, with the list of the corresponding antibiotic concentrations. Representative curves are shown from a set of at least three independent experiments. Growth delays (Δt) of treated strains relative to the growth profile without antibiotics for the *resistant* and *sCRISPRi* strains (four for meropenem assays and one for colistin assays), corresponding to the guide RNA designs of panel (a). Bars represent means and standard deviations (N=3). The bars over the interrupted axis indicate the antibiotic concentration (MIC) for which OD_600_ was lower than 0.1. **(c)** Antibiotic concentrations inhibiting bacterial growth by at least 99% (IC_99_) in agar plate assays. IC_99_ values for meropenem and colistin are shown for the strain with HSL-inducible dCas9 (induced with 10 nM HSL) and the arM1 and arM1C1 CRISPRi arrays, respectively. Bars represent the antibiotic concentrations in at least two independent experiments that provided consistent values, therefore no error bars are present. **(d)** Illustration of the experimental setup used to study escaper populations that showed recovery after antibiotic treatment in microplate assays. Supernatants of three *sCRISPRi* strains were also assayed to qualitatively investigate the antibiotic concentration remaining at the end of the experiment (Figure S15). **(e)** Analysis of escaper strains in four different conditions: arM1, arM1M2 and arM1C1 with meropenem treatment, and arM1C1 with colistin treatment. Representative curves without antibiotics and with a sub-lethal antibiotic concentration are shown for a microplate assay (first round). Three growth profiles are shown for single colonies of the escapers isolated from the treated culture of the first round and assayed as above (second round), each classified with a phenotype (resistant or resilient). Sequencing statistics are shown for each escaper, with ticks and crosses indicating correct and mutated sequences, respectively, for dCas9, CRISPRi array, and ARG. All the data come from experiments in LB media in the presence of IPTG and HSL. TOP10F’, M-res, M-cr_arM1_/M-cr_arM2_/M-cr_arM1M2_/M-cr_arM1C1_ were used as *sensitive, resistant* and *sCRISPRi* strains, respectively, to investigate meropenem re-sensitization. TOP10F’, C-res and C-cr_arM1C1_ were used as *sensitive, resistant* and *sCRISPRi* strains, respectively, to investigate colistin re-sensitization.

### Microplate assays for meropenem re-sensitization in laboratory strains

First, we examined the antibiotic response in the six strains for investigating *bla*_NDM-1_ inhibition (four *sCRISPRi* strains with gNDM1, gNDM2, gNDM1-gNDM2, gNDM1-gMCR1, and their *sensitive* and *resistant* controls). Growth delays and representative growth profiles at different MER concentrations are reported in Figure 3b and Figure S12, respectively. We observed that the *resistant* strain was able to withstand even the highest MER concentration tested, without relevant growth defects, resulting in a MIC >500 µg/ml, which was more than 500-fold higher than the one of the *sensitive* strain (MIC<1 µg/ml). In the gNDM-1 *sCRISPRi* strain, a significant growth delay is already evident for the lowest MER concentration (1 µg/ml). High delay values (up to 17 h) are reached for MER=100 µg/ml, with MIC determined at 200 µg/ml (Figure 3b), that is more than 200-fold higher than that of the *sensitive* strain. Conversely, gNDM2 caused a growth delay up to about 8 h only for much higher antibiotic concentrations (MER=500 µg/ml) with no complete growth inhibition, and statistically significant delay values only for MER>100 µg/ml (Figure 3b). This result was consistent with the previously described data on the *bla*_TEM-116_ CDS repression (Figure S4). The use of the gNDM1-gNDM2 array showed similar results to the strain with gNDM1 only, suggesting that an additional guide RNA did not improve silencing efficiency. Regarding gNDM1-gMCR1, we observed only a slight decrease in efficiency, while growth delay values comparable to gNDM1 and gNDM1-gNDM2 (less than 1.5-fold difference) and the same MIC (200 µg/ml) were determined (Figure 3b); a statistical comparison among the data of gNDM1, gNDM1-gNDM2 and gNDM1-gMCR1 for each MER concentration demonstrated that the system was not significantly affected by competition with a second spacer targeting another sequence (p>0.05, ANOVA). Together, the data on TC, AMP and MER re-sensitization showed that our one-plasmid CRISPRi platform with inducible dCas9 resulted in similar performance between individual guide RNA and two-spacer systems when targeting ARGs present in vLC and MC plasmids. This encourages the use of two-spacer CRISPRi arrays potentially targeting multiple resistance genes.

### Microplate assays for colistin re-sensitization in laboratory strains

Then, we examined antibiotic response in the three strains for investigating COL resistance repression (Figure 3b and Figure S13). We observed that the growth of the *resistant* strain was almost unaffected up to COL=2 µg/ml, and the final MIC determined (8 µg/ml) was 8-fold higher than that of the *sensitive* strain. In the gMCR-1 *sCRISPRi* strain, we determined a 2-fold lower MIC than that of *resistant* strain (Figure 3b). Moreover, comparing the growth profiles and growth delays of *sCRISPRi* and *resistant* strains, the effect of CRISPRi silencing resulted in a gradual increase of growth delay for sub-lethal COL levels, with values that were systematically higher and statistically different from the *resistant* strain at the 1 and 2 µg/ml antibiotic concentrations (p<0.05, t-test). Overall, these data showed a clear effect on colistin re-sensitization even using only one gRNA integrated in a double-spacer CRISPRi array.

### Plate assays for confirming re-sensitization to meropenem and colistin

The IC_99_ values of *ndm*- and *mcr*-*sCRISPRi* strains (0.125 µg/ml of MER and COL) were 80- and 16-fold lower than those of *resistant* controls (MER=10 µg/ml and COL=2 µg/ml) (Figure 3c). Moreover, these values approached the IC_99_ of the *sensitive* strain, showing only 4-fold difference (Table S3).

### Analysis of escapers to meropenem and colistin administration

Single colonies of *sCRISPRi* strains recovered from one of the highest MER and COL concentrations in microplate assays were collected from three independent experiments and analyzed in a second round of treatment to gain insight into the adaptation factors supporting their re-growth (Figure 3d). Indeed, the comparison of growth profiles after first and second round treatment enables the discrimination between a resistant phenotype (similar to *resistant* strain) or resilient phenotype (similar to *sCRISPRi* strain)^37^. We first studied MER-escapers and we observed both resilient and resistant phenotypes after exposure to MER (Figure 3e, Figure S14), suggesting the presence of heterogeneous populations evolving through community and adaptive antibiotic responses. Consistent with these data, the concentration of MER in the supernatant of recovered strains was lower than in the culture of a *sensitive* strain, probably due to antibiotic degradation by secreted β-lactamase, from both mutated and resilient cells, and supporting a permissive environment for resilient escapers (Figure S15). Conversely, the delay pattern of gMCR-1 *sCRISPRi* strain always resembled that of *resistant* control, as highlighted by the decreased delay in growth recovery and the increased MIC value, suggesting the selection of a population of resistant escapers (Figure 3e and Figure S14). Sequencing and restriction mapping on plasmid DNA revealed the absence of mutations in resilient cells, while inactivating mutations were found in resistant cells within the CRISPRi array (deletion of one or both spacers, 5 strains) or the dCas9 gene (insertion sequence, 3 strains). Importantly, no mutation was detected in ARG sequences, as well as no relevant copy number variation for the ARG-carrying plasmid (evaluated by electrophoresis gel analysis).

These data indicated that CRISPRi devices can re-sensitize recombinant *E. coli* strains against clinically-relevant antibiotics, although escapers with an inactivated CRISPRi system still occur at sub-lethal MER- and COL-concentrations.

## Horizontal transfer of CRISPRi repression devices in clinical isolates

We tested our CRISPRi platform in *E. coli* clinical isolates to evaluate its use for clinically relevant applications^44–46^. Notably, clinical isolates can harbor a vast collection of ARGs and genomic mutations that cooperate to generate high-level resistance^47^. In addition, the low transformation efficiency of non-model bacteria may hinder their engineering with synthetic circuits^48^. To address CRISPRi delivery, we adopted a trans-conjugative platform consisting of pTA-Mob helper plasmid, bearing the RK2 conjugative machinery^49^, and a mobilizable vector, carrying the CRISPRi circuitry and an origin of transfer (oriT from pRK2) (Figure S16a).

Platform functioning was first assessed on the previously investigated AMP-, MER- and COL-resistant laboratory strains to measure conjugation frequency and the percentage of transconjugants re-sensitized to a target antibiotic. After 24 h mating with recipients carrying *bla*_TEM-116_, *bla*_NDM-1_ or *mcr-1* gene, we determined a conjugation frequency ranging from 9*10^−5^ to 2*10^−2^ (Figure S16b), consistent with the values reported in the literature^19^. We then quantified the proportion of AMP-, MER- and COL-resistant transconjugants effectively re-sensitized to the respective target antibiotic, and we found that it ranged from 71% to 99.9% (Figure S16c).

On the one hand, these data confirmed that the conjugative transfer of CRISPRi circuits targeting three different ARGs can effectively restore the susceptibility to a fixed antibiotic concentration in almost all the population of engineered bacteria that received the mobilized plasmid; on the other hand, the low efficiency of the conjugative transfer still remains a limit of this approach, as acknowledged in the literature^50^.

### Re-sensitization of meropenem-, colistin- and cefotaxime-resistant clinical isolates

Two *E. coli* strains expressing *bla*_NDM-5_ or *mcr-1* genes were first adopted to investigate the ability of CRISPRi to increase susceptibility to the same target antibiotics as in the previous section, i.e., MER and COL. The MER-resistant isolate is a high-risk multiresistant ST167 clone collected from a liver sample of a kitten^45^. This isolate is of particular concern due to the presence of multiple β-lactamase determinants, including *bla*_ble_, *bla*_AmpH_, *bla*_AmpC1_. The COL-resistant isolate is an ST617 clone recovered from a patient admitted to the rehabilitation ward of a hospital in Northern Italy^44^. For each recipient strain, conjugations were carried out to generate transconjugants bearing a specific CRISPRi array (henceforth *sCRISPRi*^*trans*^) targeting *bla*_NDM-5_ or *mcr-1* gene, and transconjugants bearing a non-specific CRISPRi array (henceforth *nsCRISPRi*^*trans*^) used as control strains.

Growth inhibition assays were performed in Mueller Hinton (MH) broth, a nutrient-rich medium recommended by EUCAST and CLSI for antibiotic susceptibility testing. Results of microplate assays over 24 h time courses revealed that the susceptibility to both MER and COL was successfully increased in *sCRISPRi*^*trans*^ strains, exhibiting a >4- and 2-fold lower MIC than that of control strains, respectively (Figure 4b-c). Moreover, we performed MH agar plate assays and we observed that the concentration of surviving transconjugants decreased up to 163- and 4-fold for MER=1 μg/ml and COL=0.25 μg/ml, respectively (Figure S17).

**Figure 4.**
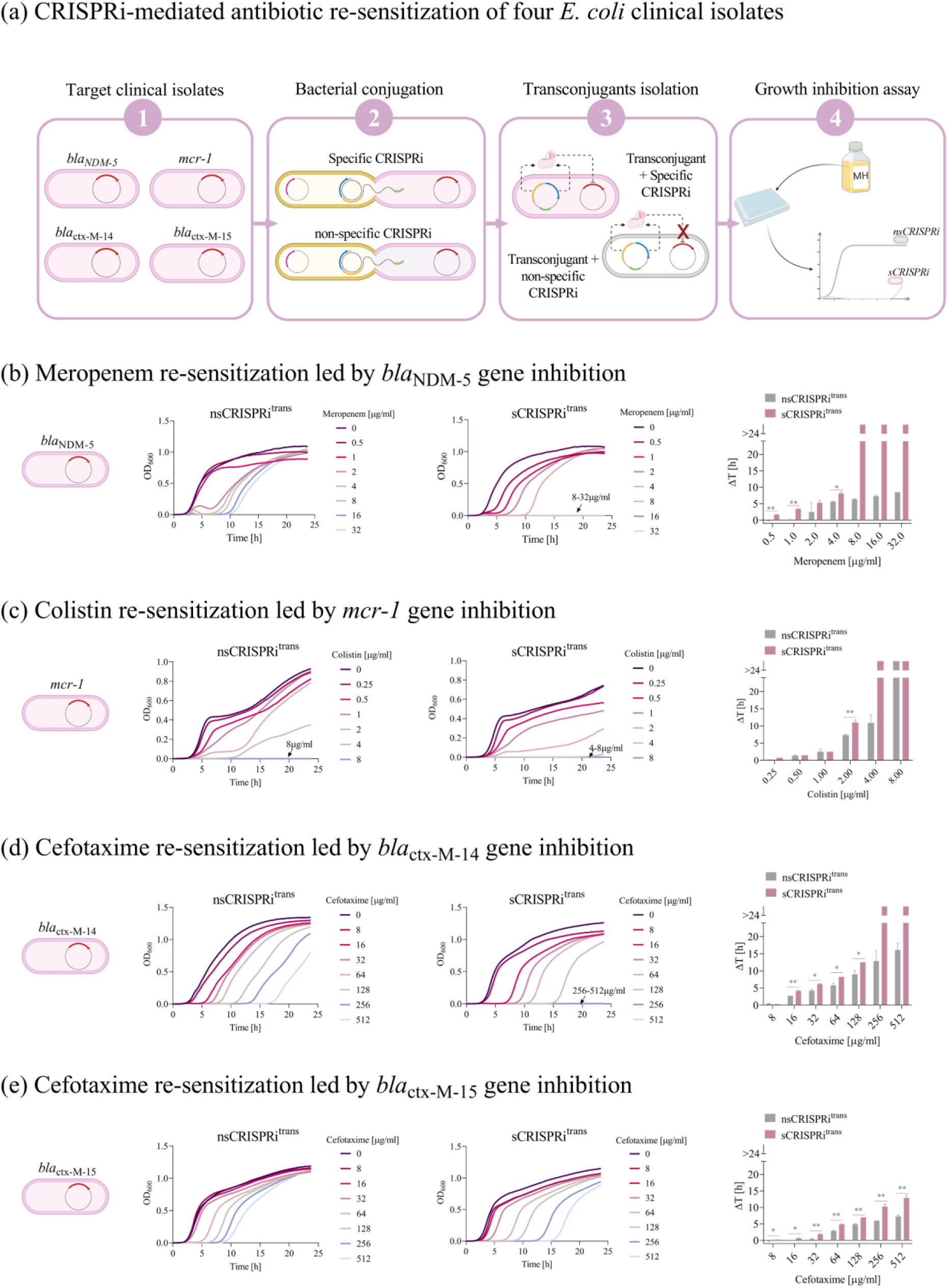
Antibiotic re-sensitization of four *E. coli* clinical isolates grown in Mueller Hinton broth. **(a)** Illustration of the experimental setup involving four clinical isolates used as CRISPRi target strains. For each isolate, two conjugations were performed to deliver a CRISPRi array transcribing a targeting or non-targeting gRNA, generating the respective transconjugants with specific (*sCRISPRi*^*trans*^) and non-specific (*nsCRISPRi*^*trans*^) CRISPRi arrays. Transconjugant strains were tested in liquid Mueller Hinton (MH) broth in microplate assays. **(b-e)** Growth profiles and delays of *nsCRISPRi*^*trans*^ (subpanel on the middle left) and *sCRISPRi*^*trans*^ (subpanels on the middle right) from microplate assays to investigate meropenem (b), colistin (c) and cefotaxime (d,e) re-sensitization. *sCRISPRi*^*trans*^ were engineered with a one-plasmid CRISPRi platform, including HSL-inducible dCas9 and IPTG-inducible CRISPRi arrays targeting the *bla*_NDM-5_ (b), *mcr-1* (c), *bla*_ctx-M-14_ (d) or *bla*_ctx-M-15_ (e) gene. The overlapped curves in which a complete growth inhibition was achieved are indicated with an arrow, with the list of the corresponding antibiotic concentrations. Representative curves are shown from a set of at least two independent experiments. Growth delays (Δt) of treated strains relative to the growth profile without antibiotics are shown for *nsCRISPRi*^*trans*^ and *sCRISPRi*^*trans*^. Bars represent the mean of two independent replicates with error bars indicating standard deviations and asterisks indicating statistical significance between the delay values of *nsCRISPRi*^*trans*^ and *sCRISPRi*^*trans*^ strains (t-test; *, p<0.05; **, p<0.01; ***, p<0.001). The bars over the interrupted axis indicate the antibiotic concentration (MIC) for which OD_600_ was lower than 0.1. All the data come from experiments in MH media in the presence of IPTG and HSL. M-cr^CI^_arAT_, C-cr^CI^_arAT_, X14-cr^CI^_arM(wt)_, X15-cr^CI^_arM(wt)_ were used as *nsCRISPRi*^*trans*^ to investigate meropenem, colistin and cefotaxime re-sensitization. M-cr^CI^_arM(wt)_, C-cr^CI^_arM1C1,_ X14-cr^CI^_arX14,_ X15-cr^CI^_arX15_ were used as *sCRISPRi*^*trans*^ to investigate meropenem, colistin and cefotaxime re-sensitization.

To further evaluate the potential of our platform in different clinical isolates and with a different antibiotic, we investigated the cefotaxime re-sensitization in two *E. coli* strains expressing the *bla*_ctx-M-14_ and *bla*_ctx-M-15_ genes. CTX-M enzymes represent a group of extended-spectrum β-lactamases (ESBLs) disseminated worldwide, with CTX-M-14 and CTX-M-15 being the most prevalent variants in gram-negative pathogens^51,52^. The two clinical isolates are part of the Natural Isolates with Low Subcultures (NILS) pathogenic *E. coli* collection^46^, and were collected from blood (*bla*_ctx-M-14_ strain) and urine (*bla*_ctx-M-15_ strain) samples. It is worth noting that, in addition to the *bla*_ctx-M-14_ gene, the same strain carries *bla*_TEM-1B_ which contributes to providing resistance to β-lactam antibiotics.

A new set of CRISPRi arrays carrying a single spacer, gCTX-M-14 or gCTX-M-15, was designed to individually target the respective ARGs (Figure 4a). Results of 24 h time courses with *bla*_ctx-M-14_ strains showed that the growth of the resistant control was not inhibited at any of the tested CTX concentrations, while a MIC was observed for the *sCRISPRi*^*trans*^ (256 µg/ml), suggesting that the CRISPRi inhibition of *bla*_ctx-M-14_ can increase by >2-fold the sensitivity to cefotaxime in MH. Consistent with these observations, a significant difference was found in the growth delays of both strains at sub-lethal CTX levels (p<0.05, t-test), with differences up to 3.5 h (Figure 4d). On the other hand, none of the CTX concentrations tested resulted in a complete growth inhibition over 24 h for *bla*_ctx-M-15_ strains, thus preventing MIC determination. However, the effect of CRISPRi on *bla*_ctx-M-15_ inhibition was still highlighted by a significant increase of growth delay values from *sCRISPRi*^*trans*^ to *nsCRISPRi*^*trans*^, with statistically significant differences for nearly all the CTX concentrations tested (p<0.05, t-test) (Figure 4e).

### Re-sensitization of meropenem-, colistin- and cefotaxime-resistant clinical isolates in LB and M9 media

Antibiotic-resistant bacteria can colonize different environments in which they adapt their growth based on available resources. Given this aspect, we tested whether the repression capability of our platform was robust under different experimental conditions affecting the growth of CRISPRi strains. In particular, we performed growth inhibition assays in rich and poor media to evaluate how the composition of these media affected the resistance profile of target strains and, accordingly, the CRISPRi repression capability. Indeed, the role played by nutrient composition in modulating bacterial regrowth kinetics and antibiotic susceptibility was previously shown^53^.

MER-, COL- and CTX-microplate assays were therefore performed in two standard growth media: minimal M9 medium with 0.1% lactose, selected as a nutrient-poor media based on inorganic salts and a carbon source, and the complex LB medium, used for assaying laboratory strains throughout this work and based on amino acids as main carbon sources, like MH (Figure 5, Figure S18).

**Figure 5.**
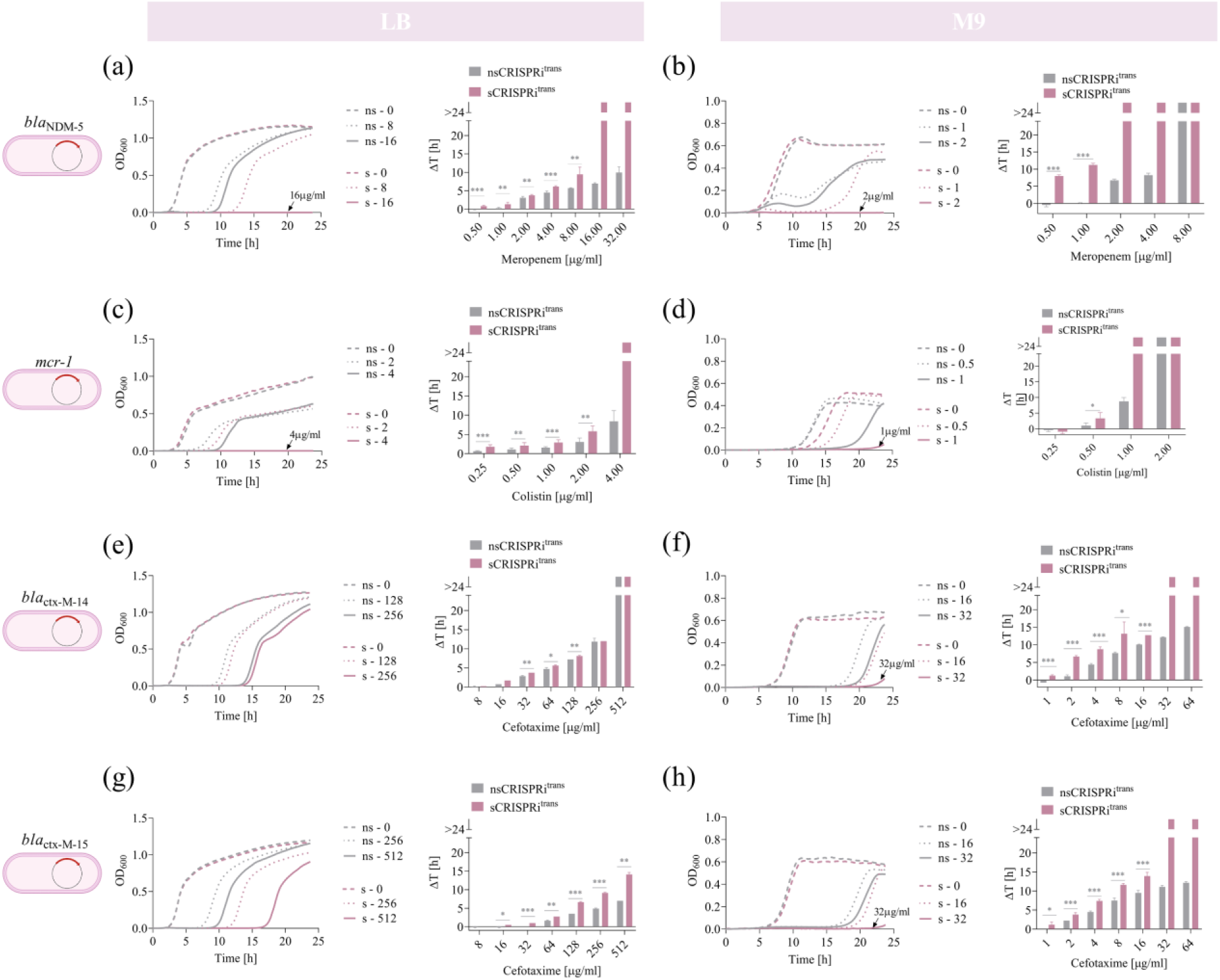
Impact of growth media conditions on antibiotic re-sensitization of four *E. coli* clinical isolates grown in L-broth and minimal M9. **(a-h)** Growth profiles and delays of *nsCRISPRi*^*trans*^ and *sCRISPRi*^*trans*^ from microplate assays to investigate meropenem (a-b), colistin (c-d) and cefotaxime (e-h) re-sensitization. *sCRISPRi*^*trans*^ are engineered with the one-plasmid CRISPRi platform described in Figure 4. Data in panels (a,c,e,g) were obtained from microplate assays in LB medium, and data in panels (b,d,f,h) were obtained from microplate assays in minimal M9 medium. Growth profiles are reported as representative curves from a set of at least three independent experiments. The growth curves of *nsCRISPRi*^*trans*^ and *sCRISPRi*^*trans*^ strains are overlapped in the same subpanels, as indicated in the legend. Only a subset of antibiotic concentrations is shown as growth profiles, while a complete set of growth curves is reported in Figure S18. Growth delays (Δt) of treated strains relative to the growth profile without antibiotics are shown for *nsCRISPRi*^*trans*^ and *sCRISPRi*^*trans*^. Bars represent the mean of at least three independent replicates with error bars indicating standard deviations and asterisks indicating statistical significance between the delay value of *nsCRISPRi*^*trans*^ and *sCRISPRi*^*trans*^ (t-test; *, p<0.05; **, p<0.01; ***, p<0.001). The bars over the interrupted axis indicate the antibiotic concentration (MIC) for which OD_600_ was lower than 0.1. Data come from experiments in LB medium in the presence of IPTG and HSL, or M9 medium with lactose in presence of HSL. Strains used to investigate meropenem, colistin and cefotaxime re-sensitization are listed in the caption of Figure 4.

Considering the growth profiles of all the strains in the three tested media (MH, M9, LB), a first major effect was a systematic increase in the antibiotic sensitivity of both non-specific and specific CRISPRi transconjugants grown in M9, in which the MIC values decreased up to 16-fold compared with those observed in MH or LB media (Figure 4-5).

Considering the *bla*_NDM-5_ *nsCRISPRi*^*trans*^ strain, all the MER concentrations tested (up to 32 µg/ml) supported growth in both MH and LB media (Figure 4b and Figure 5a). A MIC of 8 µg/ml was determined in M9, representing a >4-fold higher sensitivity than in rich media (Figure 5b). Similarly, the MIC of *sCRISPRi*^*trans*^ grown in M9 (MIC=2 µg/ml) was -4 and 8-fold lower than those determined in MH (MIC=8 µg/ml) and LB (MIC=16 µg/ml), respectively (Figure 4b and Figure 5a-b).

A comparison of MIC values between *nsCRISPRi*^*trans*^ and *sCRISPRi*^*trans*^ showed significant changes between *bla*_NDM-5_*-*strains in each tested media: the highest value was achieved in MH (FC>4), followed by M9 (FC=4) and LB (FC>2) (Figure 4b and Figure 5a-b). The contribution of the CRISPRi system also impacted antibiotic susceptibility at sub-lethal MER concentrations, in terms of growth delays. Data showed that *bla*_NDM-5_ inhibition in *sCRISPRi*^*trans*^ resulted in a statistically significant increase of growth delays than *nsCRISPRi*^*trans*^ control in all the three media for most of the MER concentrations tested (p<0.05, t-test) (Figure 4b and Figure 5a-b). Strains in M9 showed the largest recovery time for sub-lethal MER concentrations (11 h for MER=1 µg/ml), representing a much higher delay than the same strains in MH and LB (maximum delay below 4 h).

Similarly, growth inhibition data for the *mcr-1* strains showed a 4-fold lower MIC in M9 (2 µg/ml and 1 µg/ml of COL for *nsCRISPRi*^*trans*^ and *sCRISPRi*^*trans*^, respectively) than in rich MH and LB media (8 µg/ml and 4 µg/ml of COL for *nsCRISPRi*^*trans*^ and *sCRISPRi*^*trans*^, respectively) (Figure 4c and Figure 5c-d). At sub-lethal COL levels, CRISPRi increased the delay up to 2 h (M9 at COL=0.5 µg/ml) and 3 h (LB at COL=2 µg/ml) with statistically significant differences from the resistant control (p<0.05, t-test) (Figure 5b-c).

Results of growth inhibition assays with *ctx-M-*strains were consistent with previous observations: 24 h time courses performed in LB and MH media generated comparable outcomes in terms of MICs and growth delays, and all the tested strains showed a much higher increase in CTX-sensitivity when grown in M9 medium (Figure 4d-e and Figure 5e-h).

In particular, the results of *ctx-M-14* inhibition in M9 media (Figure 5f) resembled those observed in MH (Figure 4d), since a MIC (32 µg/ml) could be determined only for the strain bearing the specific CRISPRi, eventually resulting in a >2-fold improvement of CTX-sensitivity compared with the control. As in MH, the effect of CRISPRi silencing in M9 was detectable as a significant increase in the recovery time of *sCRISPRi*^*trans*^ compared with its control, with the highest difference at CTX=8 µg/ml (5.5 h) (Figure 5f). Conversely, the two *ctx-M-14* strains in LB medium displayed the same MIC (512 µg/ml) and growth delays showed differences only up to 1 h (Figure 5e). Despite the difference between both strains being statistically significant for CTX in the range 32-128 µg/ml (p<0.05, t-test), a less effective re-sensitization for *ctx-M-14* was achieved in LB than in MH and M9. It may be caused by a weaker efficiency of the CRISPRi system in this strain due to the presence of an additional non-targeted β-lactamase and/or by a faster recovery of antibiotic-inhibited survivors in this medium condition.

Finally, considering *ctx-M-15* strains, we found that the growth inhibition assays performed in LB prevent the determination of MIC values among the CTX concentration tested (Figure 5g), as already observed in MH (Figure 4e). However, CRISPRi repression on the *ctx-M-15* still promoted increased sensitivity to CTX in M9 medium, in which the MIC of *sCRISPRi*^*trans*^ was decreased by >2-fold (32 µg/ml) (Figure 5h). Moreover, growth profiles showed a significant increase of growth delay values from *nsCRISPRi*^*trans*^ to *sCRISPRi*^*trans*^, with statistically significant differences for nearly all the CTX concentrations tested (p<0.05, t-test) and values up to 5.5, 4.5 and 7 h in MH, M9 and LB, respectively (Figure 5e and Figure 5g-h).

Taken together, data on *E. coli* isolates demonstrated that our trans-conjugative CRISPRi platform could be transferred into clinically relevant pathogens, in which the sensitivity to three different antibiotics has been successfully increased. In particular, we noticed that antibiotic sensitivity increased as a function of nutrient composition of the growth media, as evidenced by the comparison between MIC values in minimal, i.e., M9, and rich media, i.e., MH and LB. However, we found that the MIC fold-changes between *nsCRISPRi*^*trans*^ and *sCRISPRi*^*trans*^ did not substantially differ in the tested media. These data demonstrated the robustness of our platform under different growth conditions since the magnitude of CRISPRi repression efficiency was conserved among three different experimental conditions. The only exception was represented by the CTX-M-15 producing strain, in which the contribution of CRISPRi in decreasing the MIC was only detectable in M9 medium. In addition to MIC determination, the analysis of the growth delays of the tested strains treated with sub-inhibitory antibiotic concentrations allowed us to enrich our understanding of the effective potential of the CRISPRi system, which was reflected in a statistically significant increase of the recovery time for *sCRISPRi*^*trans*^ strains with respect to their controls in almost all the tested conditions. Apart from the different antibiotic sensitivity, the media-dependent behavior showed different patterns among the tested strains, with one of them showing no biologically relevant CRISPRi effectiveness in one of the three tested media, and another strain failing to reach a MIC value.

## Discussion

The reversion of antibiotic resistance is one of the innovative approaches conceived to counteract the emergence of multidrug-resistant pathogens and the slow pace in developing new antimicrobial drugs. CRISPR-based tools have been previously proposed for sequence-specific targeting of antibiotic-resistant pathogens, to carry out double-strand DNA breaks (DSB) that can kill the strain or re-sensitize it to drugs that it no longer responds to. However, some challenges still need to be addressed to properly evaluate the translation of this technology into medical settings. Examples are the *in-vivo* transfer rate of CRISPR circuits and the risk of target gene mutations driven by the SOS response. In this work, we proposed the use of a catalytically-inactive Cas9 (CRISPRi) to re-sensitize target strains via transcriptional inhibition of resistance genes, accordingly minimizing the risk of generating new resistant variants. We tested the effectiveness of CRISPRi under challenging conditions never tested in previous studies, including the inhibition of one or more ARGs placed on low- to high-copy plasmid, or in the bacterial chromosome of *E. coli* laboratory strains and different clinical isolates.

A preliminary design of our genetic platform relied on two plasmids individually carrying the expression cassettes of gRNA and dCas9, which was expressed via a constitutive promoter. Using this configuration, we targeted the promoter and/or coding sequence of tetracycline- and ampicillin-resistance genes in *E. coli* laboratory strains. In these tests, we demonstrated an effective re-sensitization to both antibiotics, since we determined a decrease of MIC values and colony counts in liquid and solid media assays. Furthermore, this configuration enabled ampicillin re-sensitization even if the correspondent ARG was placed in a high copy plasmid.

The genetic platform design was then optimized 1) to place dCas9 and guide RNA in the same plasmid, 2) to enable the targeting of multiple ARGs, via dCas9 expression tuning, and 3) to compare different architectures for guide RNA expression (sgRNA tandem cassettes and CRISPRi arrays). We observed that the simultaneous repression of tetracycline- and ampicillin-resistance genes required a proper tuning of dCas9 expression level. Moreover, when comparing the two gRNAs architectures, we observed a slightly higher repression performance for the CRISPRi array, which was eventually selected for our study. We reprogrammed the optimized platform to test the re-sensitization to last-resort drugs, i.e., meropenem and colistin, by using CRISPRi arrays with one or two spacers. Overall, these four case studies in laboratory strains demonstrated that our platform enables the repression of two ARGs with efficiency comparable to one-guide systems when the target is present in low- to medium-copy plasmids.

We also analyzed escapers recovering from experiments in microplate assays to gain insight into their escape mechanism. We found that both mutations breaking the CRISPRi circuitry and/or the emergence of community-dependent behavior (e.g., β-lactams inactivation due to collective antibiotic tolerance) supported the regrowth of escapers at sub-MIC antibiotic treatments. However, we did not find mutations in the target gene, as observed with CRISPR antimicrobials.

Finally, to test the re-sensitization capability of our platform in more relevant therapeutic contexts, we delivered the CRISPRi plasmid into four *E. coli* clinical isolates via bacterial conjugation. We selected these strains due to the presence of ARGs of severe clinical concern (e.g., *bla*_NDM-5_ and *mcr-1*) and other additional genes/mutations that may work together to provide a high resistance level. We tested the robustness of the platform by performing growth inhibition assays with meropenem, colistin and cefotaxime (two ARG variants) under different growth conditions (MH, LB, M9 media). We observed a strong medium-dependent sensitivity when determining the MIC of both specific CRISPRi and control strains with a non-targeting system. Conversely, the fold increase of their MIC between specific and non-specific CRISPRi strains was more conserved. In particular, we observed that in standard M9 the sensitivity to the three antibiotics was increased by 2- to >4-fold in the tested strains. However, re-sensitization capability has been found to be more variable in MH and LB.

These data confirm the complex interplay between genetic and environmental factors contributing to antibiotic resistance in bacteria. In this direction, our platform could provide a tool to dissect AMR by perturbing the expression of specific genes (PBPs, porins, efflux pumps, etc.) individually or in combination.

Together, we demonstrated that CRISPRi can be a promising re-sensitization tool against last resort antibiotics in high-risk multidrug-resistant clinical isolates. Further platform design efforts will include different markers and/or regulatory elements to enable the selection and expression of CRISPRi complex in target strains. Although some limitations still exist in applying this technology as a viable therapeutic, the work herein presented laid the foundations for further investigating the potential of CRISPRi in the global race against antibiotic resistance.

## Materials and Methods

### Strains and plasmids

The strains used in this work are listed in Table S1, while their plasmids and scope are detailed in Table S2. Most of them are derivatives of the TOP10F’ parent strain with the F’ episome conferring tetracycline resistance by the *tetA* gene, and overexpressing the LacI repressor via the *lacI*^*q*^ locus. Exceptions are: the TOP10 strain with no F’ plasmid that was used as a tetracycline sensitive control, the DH10B strain that was used as a donor chassis for conjugation assays, and the *E. coli* clinical isolates that were used as recipient strains^44–46^. All the strains were long-term stored at -80°C in 20% glycerol stocks.

The pI13521 plasmid^35^, based on the pSB1A2 backbone with the mutated pMB1 origin conferring high-copy vector maintenance per cell, was used as the ampicillin resistance plasmid. It constitutively expresses *bla*_TEM-116_, coding for a β-lactamase, by the P_bla_ promoter.

The pJ107202 plasmid^35^, based on the medium-copy kanamycin-resistant pSB3K3 backbone with the p15A origin, was used as a constitutive source of dCas9, via an expression cassette driven by the weak P_J23116_ promoter.

The pGDP1 NDM-1 and pGDP2 MCR-1^43^ were gifts from Gerard Wright (Addgene plasmids #112883 and #118404). They were used as constitutive expression vectors for a codon-optimized *bla*_NDM-1_ (by the P_bla_ promoter) and *mcr-1* (by the P_lac_ promoter), respectively. Both plasmids have a kanamycin resistance gene and are replicated at medium-copy number via the pBR322 origin.

The pTA-mob plasmid^49^ was a gift from Rahmi Lale (NTNU). This helper plasmid includes a gentamicin resistance marker, a pBBR1 broad-host range replicon conferring medium-copy maintenance, and provides *in trans* all the genes necessary for conjugative transfer of mobilizable plasmids that have an origin of transfer (oriT).

The chloramphenicol-resistant low-copy pSB4C5 backbone^35^, having the pSC101 origin, was adopted for cloning, heterologous expression of synthetic circuits (P_LlacO1_-driven sgRNAs or CRISPRi arrays, P_lux_-driven dCas9 in HSL-inducible dCas9 circuits), and as a mobilizable plasmid using the oriT sequence, amplified from the RK2 plasmid (ATCC 37125)^54^.

### Reagents and media

Isopropyl-β-D-1-thiogalactopyranoside (IPTG, #I1284, Sigma Aldrich) and N-3-oxohexanoyl-L-homoserine lactone (HSL, #K3007, Sigma Aldrich), routinely stored at -20°C, were used as chemical inducers of recombinant gene expression for RNA guides (sgRNAs or CRISPRi arrays) and for dCas9 in the circuit architectures with inducible dCas9, respectively. Ampicillin (AMP, 100 mg/ml), kanamycin (KAN, 50 mg/ml), chloramphenicol (CHL, 34 mg/ml), gentamicin (GM, 50 mg/ml), tetracycline (TC, 5 mg/ml) stocks were routinely stored at -20°C. The powders of levofloxacin (LEV), ciprofloxacin (CIP), meropenem (MER), colistin (COL) and cefotaxime (CTX) were stored at -20°C and used to make fresh stocks at every use. Antibiotic powders were dissolved in deionized water except chloramphenicol (ethanol) and meropenem (deionized water with 10.4 mg/ml of sodium carbonate). All of them were filter-sterilized (0.2 μm) before storage. Meropenem and colistin agar plates were used within two days from preparation.

Unless differently indicated, L-broth (LB; 10 g/L tryptone, 5 g/L yeast extract, 10 g/L sodium chloride) was used as a medium for cloning and quantitative experiments. Agar plates were prepared by adding 15 g/L agar to the liquid medium before autoclaving. Mueller Hinton (MH) and M9 (M9 salts 11.28 g/L, MgSO_4_ 2 mM, CaCl_2_ 0.1 mM, and 0.1% lactose as carbon source) were used as liquid media in the experiments with clinical isolates. Tween 20 0.1% was added to M9 medium in experiments with the X14-cr^CI^_arM(wt)_, X14-cr^CI^_arX14_, X15-cr^CI^_arM(wt)_, and X15-cr^CI^_arX15_ strains (Table S1).

Selective media refers to medium only including the antibiotics needed for plasmid maintenance or strain selection (without the target antibiotic tested for growth inhibition): chloramphenicol (12.5 μg/ml), kanamycin (25 μg/ml), gentamicin (20 μg/ml), levofloxacin (2 μg/ml) and ciprofloxacin (2 μg/ml), according to the specific plasmids. Antibiotics tested for growth inhibition were ampicillin, tetracycline, meropenem, colistin and cefotaxime, added at the indicated concentrations.

### Circuit assembly

Unless differently indicated, the BioBrick Standard Assembly procedure^55^ was adopted for plasmid construction, enabling binary assembly steps that left a standard DNA junction (TACTAGAG) between the joint parts. The dCas9 expression systems and the sgRNA scaffold, with synthetic tetraloop for the expression of the gRNA:tracrRNA as a unique molecule were obtained previously^35^, the oriT sequence was PCR-amplified from RK2 and all the CRISPRi arrays were obtained by *de novo* DNA synthesis (GenScript, Piscataway, NJ, USA). Digestions with *EcoRI* and *SpeI* restriction sites in DNA parts with dCas9 and CRISPRi array, respectively, were avoided, due to illegal restriction sites in their sequences but their assembly was still possible with BioBrick restriction enzymes (*XbaI* and *PstI*). sgRNA cassettes were customized by changing the 20-nt targeting region via mutagenesis with divergent primers, using PCR amplification, *DpnI* digestion, T4 polynucleotide kinase and ligase reactions, as previously carried out^35^. Targeting regions were designed via the Benchling CRISPR tool (*https://benchling.com*) by setting a 20-nt (for sgRNAs) or 30-nt (for CRISPRi arrays) guide length, GCA_00005845.2 reference genome, and the optimized score by Doench et al.^56^. Each sgRNA cassette was designed with its own IPTG-inducible P_LlacO1_ promoter. DNA purification kits (Macherey-Nagel), restriction enzymes, T4 DNA ligase and T4 polynucleotide kinase (Thermo Scientific), Antarctic phosphatase (New England BioLabs), and Phusion Hot Start II PCR kit (Thermo Scientific) were used according to manufacturer’s indications. Sequencing and oligonucleotides synthesis services were from Eurofins Genomics Germany GmbH.

### Growth inhibition assays

Strains from long-term stocks were streaked on an LB agar plate and incubated overnight (14-16 h) at 37°C. For tests with laboratory strains, single colonies (N=3) were used to inoculate 0.5 ml of selective media in 2-ml tubes and cultures were grown overnight at 37°C, 220 rpm in an orbital shaker. The grown cultures were 100-fold diluted in 200 μl of selective LB in a 96-well microplate, in which we added IPTG (500 μM) and HSL (10 nM, unless differently indicated) to trigger guide RNAs and dCas9 expression, as required, and antibiotics were added to reach the desired concentrations. The 96-well plate was incubated at 37°C in an Infinite F200Pro reader (Tecan) and assayed for 24 h via an automated sampling program (i-control software, Tecan): linear shaking (5 s, 3-mm amplitude), 5 s wait, absorbance measurement (600 nm), 5-min sampling time.

For tests involving clinical isolates, strains were grown as previously described^57^ to start experiments with exponentially growing cells at comparable densities. Briefly, colonies were used to inoculate 2 ml of selective media in 50-ml tubes and cultures (N=3) were grown overnight at 37°C, 200 rpm in an orbital shaker. The overnight cultures were diluted to 0.05 OD_600_ in fresh media and incubated for 3-4 h under the same conditions as above. These cultures were finally diluted to 0.0005 OD_600_ in 200 μl of the indicated medium in a 96-well microplate, in which IPTG (500 μM), HSL (10 nM), and the required antibiotics were added to reach the desired concentrations. IPTG was omitted when using M9 media with lactose, which already activates the P_LlacO1_ promoter. The microplate was assayed as previously described^57^ at 37°C in a Spark reader (Tecan) for 24 h via the following kinetic cycle programmed with Spark Control v.2.2 software (Tecan): linear shaking (30 s, 5 mm), orbital shaking (30 s, 5 mm), 30 s wait, absorbance measurement (600 nm), 60-s wait, linear shaking (120 s, 5 mm), 30 s wait, orbital shaking (120 s, 5 mm), 30 s wait, orbital shaking (60 s, 5 mm), 15-min sampling time. Sterile media without bacteria was also included to measure the absorbance background.

During the assays, antibiotics (CHL and KAN, when required) were added to solid or liquid media for the selection of strains with the CRISPRi system. No other antibiotics were added for the maintenance of ARG-bearing plasmids, for which no loss was observed during strain propagation (data not shown).

Occasionally, at the end of experiments with ampicillin and meropenem, 3 μl of bacteria-free supernatants from cultures grown in different wells were applied on selective LB agar plates on which an AMP- or MER-sensitive strain was previously plated. The plates were incubated overnight at 37°C and inhibition halos were observed, qualitatively indicating antibiotic activity and, consequently, its degradation due to the β-lactamase enzymes released in the media. Growth inhibition assays on solid media were carried out by plating serial dilutions of the recombinant cultures (N=2) grown overnight in the 2-ml tubes. LB and MH plates were supplemented with the antibiotics required for ARG-bearing plasmid selection, IPTG (500 μM) and HSL (10 nM, unless differently stated) as required, and the target antibiotic (ampicillin, tetracycline, meropenem or colistin) at different concentrations. Plates were incubated at 37°C overnight.

### Escaper analysis

At the end of the 24-h growth assays in 96-well microplates, escaper cultures recovering from one of the highest antibiotic concentrations were streaked on selective LB agar without target antibiotic and incubated overnight at 37°C. From each plate, one colony was used to inoculate 5 ml of selective LB, incubated overnight at 37°C, 220 rpm. The grown cultures were 100-fold diluted in a microplate to run a second-round microplate assay under the same conditions as the first round microplate assay that originated the escaper culture. The rest of the culture was used to extract plasmid DNA. Plasmids were used to identify mutations by sequencing with primers covering guide RNAs, dCas9 and ARG target site (see Table S3), and using restriction mapping on 1% agarose gel electrophoresis.

### Conjugation

Donor and recipient strains from a streaked agar plate were used to inoculate 5 ml of selective media and grown overnight at 37°C, 220 rpm. Cultures were 100-fold diluted and incubated under the same conditions until they reached 0.25 OD_600_. Then, 1 ml was withdrawn from each tube, gently spun down (3000 rpm, 5 min) and washed twice with sterile PBS to remove residual antibiotics from the cultures. OD_600_ was again measured and each culture was concentrated to reach a final OD_600_ of 0.5. Mating pairs were then combined in a 1:1 donor:recipient ratio and 100 μl of the mixture was pipetted on a non-selective plate that was further incubated at 37°C for 20 h. The conjugation mixture was scraped up from the plate and transferred into 1 ml PBS in a 1.5-ml tube. Conjugation was interrupted by vortexing, centrifuging (5000 rpm, 3 min) and resuspending the mating mixture in PBS, which was finally diluted as appropriate. Protocol definition was based on a screening of different parameters that maximized conjugation frequency, i.e., bacterial density, growth phase, solid vs. liquid media, strains ratio, and size of the spotted mixture (data not shown).

For conjugation of the M-res, C-res, M-res^CI^, C-res^CI^ strains, the mixture was plated on agar media (LB for M-res and C-res; MH for M-res^CI^ andC-res^CI^) with CHL + auxiliary antibiotic (i.e., KAN, LEV or CIP, as required to select ARG-bearing plasmids - see Table S1) to select for transconjugants; the same mixture was plated on media to select for recipients (auxiliary antibiotic), and survivors (CHL + IPTG/HSL + target antibiotic, i.e., MER or COL). For conjugation of A-res, having no auxiliary antibiotic in the ARG-bearing plasmid, the mixture was plated on LB agar with CHL + AMP to select for transconjugants, AMP to select for recipients, and CHL + AMP + IPTG/HSL to select for survivors to antibiotic treatment. Absence of *bla*_TEM-116_ inhibition was observed when dCas9 and sgRNA were not expressed, based on colony counts with and without AMP (data not shown).

For conjugation of the X14-res^CI^, X15-res^CI^ strains, for which no auxiliary antibiotics were exploited, the mixture was plated on minimal M9 plates with 0.4% lactose with CHL to select for transconjugants (the DH10B-based donor was unable to grow in minimal media without leucine and was also unable to consume lactose), and on the same agar media without CHL to select for recipients.

In all cases, plates were incubated overnight at 37°C and the colony forming units (CFUs) for transconjugants, recipients and survivors were quantified as *N*_*T*_, *N*_*R*_ and *N*_*S*_, respectively. These CFU numbers were used to calculate conjugation efficiency (*E*_*conj*_), as *N*_*T*_*/N*_*R*_, and the proportion of re-sensitized transconjugants, as *1-N*_*S*_*/N*_*T*_. Unless differently indicated, three independent experiments were performed for each condition.

## Data analysis

For each sample in microplate assays, raw absorbance time series was background-subtracted using the raw measurement of sterile media, obtaining an OD_600_ time series that is proportional to bacterial density. The linearity of the plate reader was assessed up to OD_600_ values of 0.7.

The minimum inhibitory concentration (MIC) was determined in microplate assays as the antibiotic concentration for which cell growth is maintained lower than OD_600_ 0.1 over 24h.

Growth profiles of strains at different antibiotic concentrations were used to quantify the CRISPRi-mediated antibiotic re-sensitization by calculating Δt=t_0.1,AB+_–t_0.1,AB-_, where t_0.1,AB+_ and t_0.1,AB-_ are the time points in which the cultures with and without the tested antibiotic reach OD_600_=0.1. This index was computed for each antibiotic concentration within a biological replicate, for which the initial density in the microplate is expected to be the same since the wells were inoculated with the same pre-culture.

IC_99_ was determined in agar plate assays as the antibiotic concentration inhibiting 99% of the bacterial growth, measured as colony count, compared with the plate in the absence of the target antibiotic.

Statistical tests were carried out using Matlab R2017b (MathWorks) or Microsoft Excel to compare mean values of growth delay across different strains for each antibiotic concentration tested. Unpaired one-sided t-test was used to compare resistant and CRISPRi strains. Two-way ANOVA with replicates was used to compare circuit architectures, with dCas9 (constitutive vs. inducible) and guide RNA (double sgRNA vs. CRISPRi array) as main factors. One-way ANOVA was used to compare circuit architectures including gNDM1. A p-value of 0.05 was used as a cutoff for statistical significance. Graphs were generated using GraphPad Prism 8.3.0.

## Supporting Information

Supplementary figures: Characterization methods for quantifying the CRISPRi-mediated re-sensitization to target antibiotics (Figure S1); Growth curves of the *sensitive* strains in the presence of tetracycline or ampicillin (Figue S2); Re-sensitization to tetracycline and ampicillin in laboratory strains using a CRISPRi platform, including a weak constitutive dCas9 and an IPTG-inducible sgRNA, without IPTG-induced sgRNA expression (Figure S3); Re-sensitization to ampicillin in laboratory strain using a CRISPRi platform, including a weak constitutive dCas9 and an IPTG-inducible sgRNA, using a guide targeting the *bla*_TEM-116_ coding sequence (Figure S4); Comparison of growth delays between resistant laboratory strain and resistant laboratory strain bearing a non-specific guide RNA (Figure S5); Growth profiles of escaper mutants isolated from tetracycline and ampicillin microplate assays with laboratory strains (Figure S6); Agar plate assays for the characterization of CRISPRi-mediated single-ARG inhibition (S7); Comparison of growth delays among single-sgRNA, double-sgRNA, and CRISPRi array designs, with a weak constitutive dCas9 expression cassette in laboratory strains (Figure S8); Growth delays of laboratory strains bearing a one-plasmid CRISPRi platform, including an inducible dCas9 and a two guide RNAs architecture (double sgRNA cassette or CRISPRi array) for different HSL concentrations (Figure S9); Growth delays of the laboratory strain bearing a one-plasmid CRISPRi platform including inducible dCas9 and CRISPRi array targeting tetracycline and ampicillin resistance genes, in a combined treatment with the two antibiotics (Figure S10); Target sites of the designed gNDM1 and gNDM2 guides in the *bla*_NDM-1_ coding sequence (Figure 11); Re-sensitization to meropenem in laboratory strains using a one-plasmid CRISPRi platform, including HSL-inducible dCas9 and one of four different CRISPRi arrays targeting *bla*_NDM-1_ (Figure S12); Re-sensitization to colistin in laboratory strains using a one-plasmid CRISPRi platform, including HSL-inducible dCas9 and CRISPRi array targeting *mcr-1* (Figure S13); Growth delays of escaper mutants during second-round experiments (Figure S14); Supernatant assay with *bla*_NDM-1_-expressing laboratory strains (Figure S15); Horizontal transfer of the CRISPRi plasmid from a donor strain to antibiotic-resistant laboratory strains (Figure S16); Meropenem and colistin re-sensitization in agar plate assays for two clinical isolates with *bla*_NDM-5_ and *mcr-1* (Figure S17); Antibiotic re-sensitization of four *E. coli* clinical isolates grown in LB and M9 media. (Figure S18). Supplementary tables: List of strains used in this work (Table S1); List of recombinant plasmids used in this work (Table S2); List of primers used in this work for escaper analysis (Table S3); List of IC_99_ values determined with agar plate assays (Table S4).

## Author Contribution

A.F.C., P.M. and L.P. conceived the study. A.F.C, A.P., R.M., G.B., P.M. and L.P. designed the experiments. A.F.C. constructed the plasmids. A.F.C. and M.C. conducted the quantitative experiments. A.F.C., M.B., G.B., P.M. and L.P. analyzed the data. A.F.C. and L.P. wrote the manuscript. M.B., G.B. and P.M. contributed to manuscript finalization. All authors read and approved the manuscript.

## Data availability

The data that support the findings of this study are available from the corresponding author upon request.

## Notes

The authors declare no competing financial interest.

## Acknowledgements

The authors want to thank Daniele Pastorelli, Melissa Spalla, Aseel AbuAlshaar, Francesca Piscopiello (Univ. Pavia) and Viktoriia Gross (INRIA/Pasteur) for their help in experimental work with clinical isolates; Angelo Serafini, Mirko Martini and Marianna Caldarulo (Univ. Pavia) for their work with laboratory strains and the preliminary work on protocol design; Maria Gabriella Cusella De Angelis (Univ. Pavia) for sharing lab space and instruments. This work was supported by Innovation HUB Regione Lombardia, Italy (grant 1139857), by Fondazione Cariparo (grant 59576, 2021) and by ANR, France (grant 20-PAMR-0010, *Seq2DiAg*). Lorenzo Pasotti also acknowledges funding support from the INCEPTION visiting scientist fellowship (Institut Pasteur, ANR, Investissements d’Avenir, France).

