## Supplementary Information for "Harnessing CRISPR interference to re-sensitize laboratory strains and clinical isolates to last resort antibiotics"

### Supplementary Figures

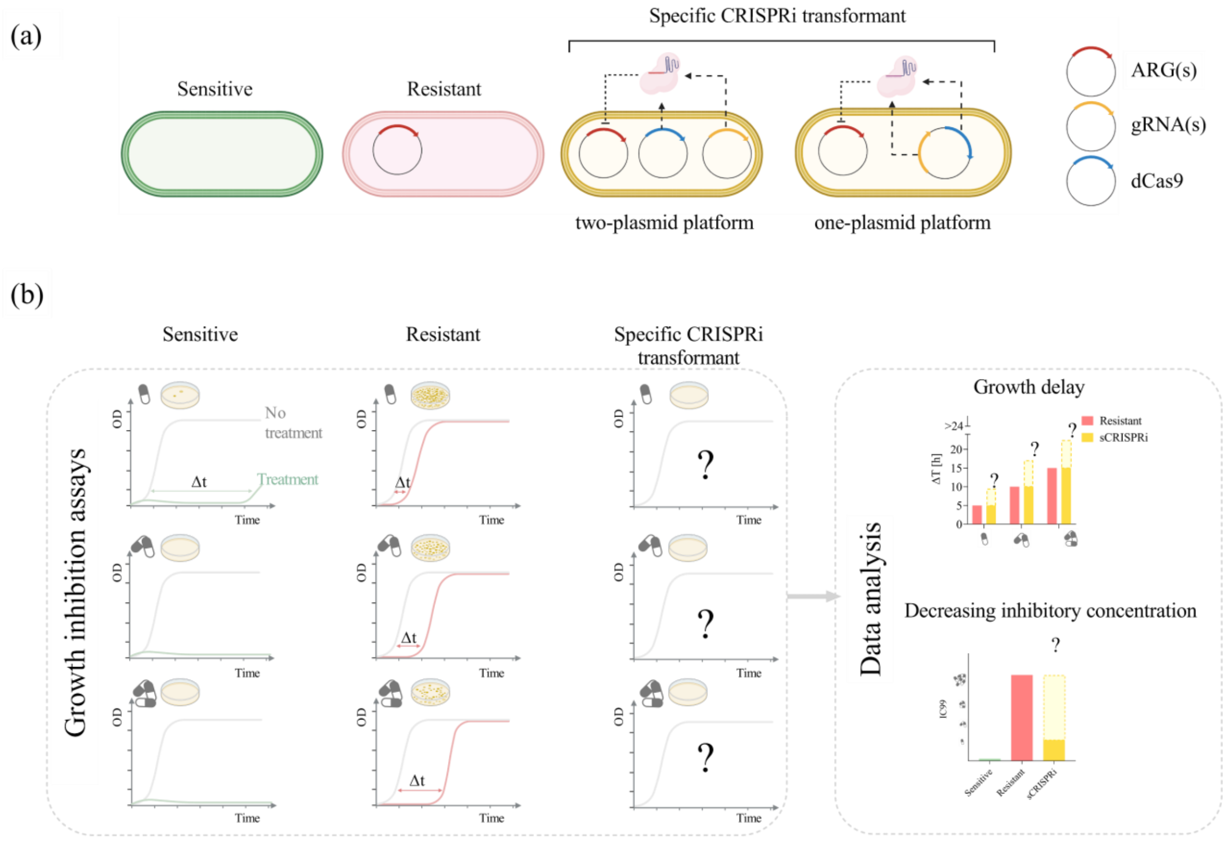

**Figure S1.**

**Characterization methods for quantifying the CRISPRi-mediated re-sensitization to target antibiotics. (a)** Description of the test strains: *sensitive*, lacking the ARG; *resistant*, bearing the ARG; *specific CRISPRi transformant* (*sCRISPRi*), bearing the ARG and the CRISPRi platform. Plasmids with their main components: ARG (red), dCas9 module (blue); and sgRNA module (yellow). The ARG(s) can be located on one plasmid, two plasmids or on the chromosome, depending on the considered case study. The dCas9 module can be driven by a weak constitutive promoter in a medium copy plasmid or by an HSL-inducible promoter in a low copy plasmid. The gRNA module has been implemented to transcribe one or two sgRNAs under an IPTG-inducible promoter in a low-copy plasmid. A constitutive dCas9 module and an IPTG-inducible sgRNA module constitute a two-plasmid CRISPRi platform. An HSL-inducible dCas9 module and an IPTG-inducible sgRNA module constitute a one-plasmid CRISPRi platform. **(b)** CRISPRi-mediated re-sensitization is quantified via growth inhibition assays. Different concentrations of target antibiotics are added in liquid media to perform microplate assays (on the left), or solid media, to perform agar plate assays (on the right). The expected phenotypes are illustrated as a growth profile in response to antibiotic treatment (illustrated as pills) with the relative growth delays from microplate assays, and $IC_{99}$ from agar plate assays.

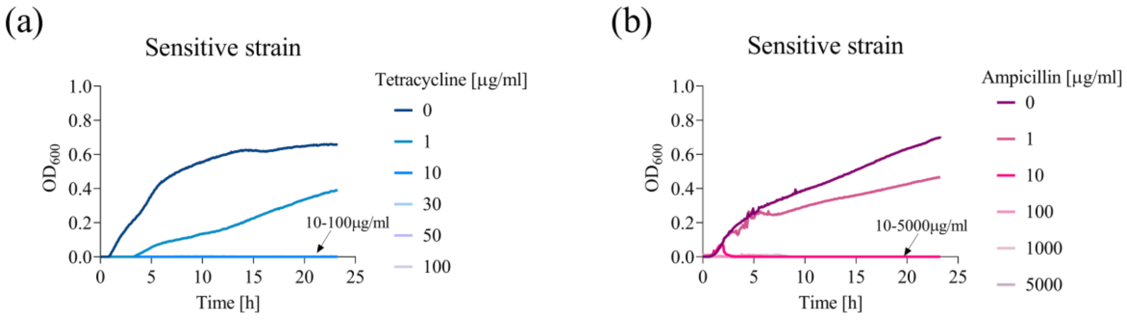

**Figure S2.**

**Growth curves of the *sensitive* strains in the presence of tetracycline or ampicillin. (a-b)** Representative curves are shown from a set of at least three independent experiments with *sensitive* strains treated with tetracycline (a) or ampicillin (b). Data come from experiments in LB media with IPTG. TOP10 and TOP10F' were used as *sensitive* strains for tetracycline and ampicillin, respectively.

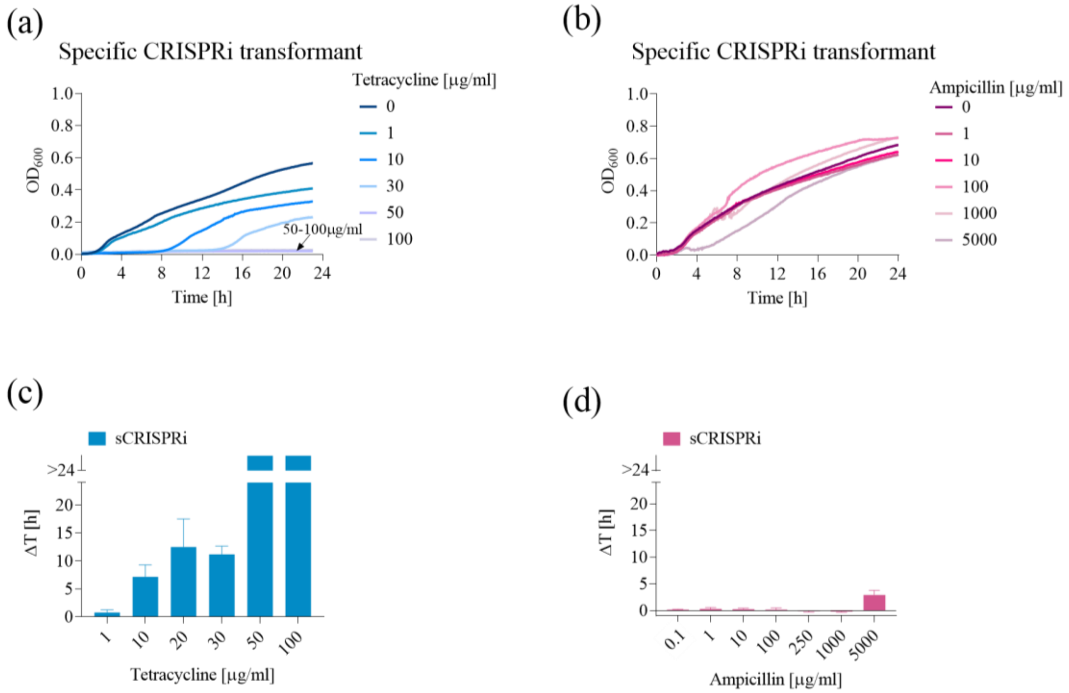

**Figure S3.**

**Re-sensitization of laboratory strains to tetracycline and ampicillin by single ARG targeting, in the absence of IPTG. (a-b)** Growth profiles of *sCRISPRi* strains from microplate assays to investigate tetracycline (a) and ampicillin (b) re-sensitization. *sCRISPRi* strains are engineered with a CRISPRi platform, including constitutive dCas9 and IPTG-inducible gtetA or $gbla_{promoter}$ targeting *tetA* or $bla_{TEM\text{-}116}$, respectively. The overlapped curves in which a complete growth inhibition was achieved are indicated with an arrow, with the list of the corresponding antibiotic concentrations. Representative curves are shown from a set of at least three independent experiments. **(c-d)** Growth delays (Δt) of treated strains relative to the growth profile without antibiotics for the *resistant* and *sCRISPRi* strains. Bars represent means and standard deviations (N=3). The bars over the interrupted axis indicate the antibiotic concentration (MIC) for which $OD_{600}$ was lower than 0.1. Different from Figure 1 in the main text, all the data presented here have been measured without IPTG that triggers sgRNA expression. $T\text{-}cr_{sgT\_const}$ and $A\text{-}cr_{sgA\_const}$ were used as *sCRISPRi* strains to investigate tetracycline and ampicillin re-sensitization, respectively.

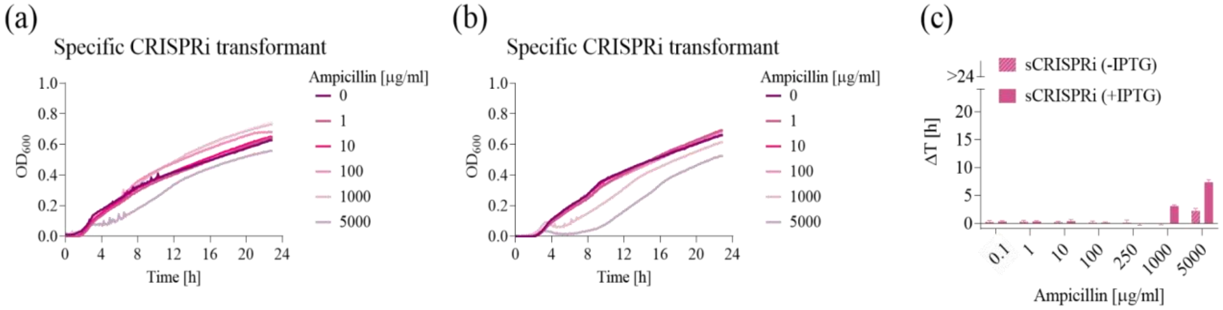

**Figure S4.**

**Re-sensitization of laboratory strains to ampicillin by targeting the *bla*$_{TEM-116}$ coding sequence, in the absence or presence of IPTG. (a-b)** Growth profiles of *sCRISPRi* strain from microplate assays to investigate ampicillin re-sensitization. *sCRISPRi* strain is engineered with a two-plasmid CRISPRi platform, including constitutive dCas9 and IPTG-inducible gbla$_{CDS}$ targeting *bla*$_{TEM-116}$ CDS. Antibiotic concentrations are provided in the legend. Representative curves are shown from a set of at least three independent experiments. Growth profiles are shown without IPTG (a) and with IPTG (b) that induced gbla$_{CDS}$ expression. **(c)** Growth delays ($\Delta$t) of treated strains relative to the growth profile without antibiotics for the *resistant* and *sCRISPRi* strains, with/without IPTG as indicated in the legend. Bars represent means and standard deviations (N=3). The bars over the interrupted axis indicate the antibiotic concentration (MIC) for which OD$_{600}$ was lower than 0.1. All the data come from experiments in LB media. A-cr$_{sgA(CDS)_const}$ was used as *sCRISPRi* strain to investigate ampicillin re-sensitization.

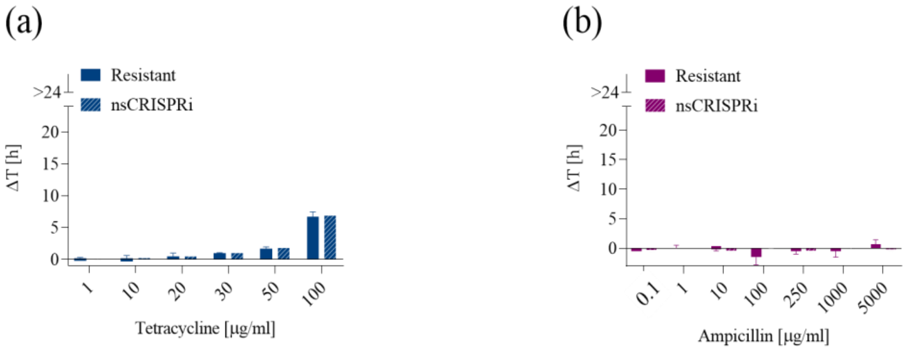

**Figure S5.**

**Comparison of growth delays between resistant laboratory strain and resistant laboratory strain bearing a non-specific guide RNA. (a-b)** Growth delays ($\Delta$t) of tetracycline (a) and ampicillin (b) treated strains relative to the growth profile without antibiotics for the *resistant* and *non-specific CRISPRi transformant* (*nsCRISPRi*) strains. Bars represent means and standard deviations (N=3). All the data come from experiments in LB media in the presence of IPTG. TOP10F' and T-cr$_{sgA_const}$ were used as *resistant* and *nsCRISPRi* strains to investigate tetracycline re-sensitization, respectively. A-res and A-cr$_{sgT_const}$ were used as *resistant* and *nsCRISPRi* strains to investigate ampicillin re-sensitization, respectively.

### Supplementary notes for Figure S6.

*Phenotypic analysis of the escapers to tetracycline and ampicillin treatment*. Since we observed the appearance of an escaper population in microplate assays, we investigated whether its origin was related with the evolution of mutations. Single colonies of *sCRISPRi* strains that recovered from the highest AMP and TC concentration in microplate assays were collected and used to carry out a second round of treatment with the same range of antibiotic concentrations. All the tested colonies (N=3) exhibited a delay pattern comparable with the *resistant* strain, as evidenced by the decreased recovery time and the increased MIC values, suggesting that the isolated escapers were represented by mutants.

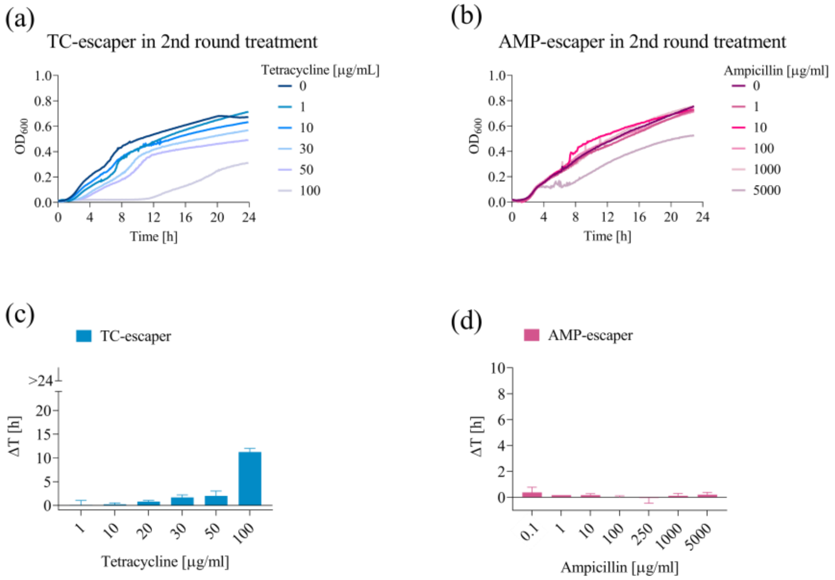

**Figure S6.**

**Growth profiles of escaper mutants isolated from tetracycline and ampicillin microplate assays with laboratory strains. (a-b)** Growth profiles of tetracycline- (a) and ampicillin- (b) escaper mutants from second round antibiotic treatment in microplate assays. Escapers were isolated at the end of 24 h time courses (first round treatment) performed with *sCRISPRi* strains bearing gtetA and $gbla_{promoter}$ and treated with one of the highest sub-lethal antibiotic concentrations. Escapers from first round treatment were treated in the second-round treatment with the antibiotic concentrations provided in the legend. Representative curves are shown from a set of at least three independent colonies. **(c-d)** Growth delays (Δt) are shown for the same strains relative to the growth profile without antibiotics. Bars represent means and standard deviations (N=3). All the data come from experiments in LB media in the presence of IPTG.

.

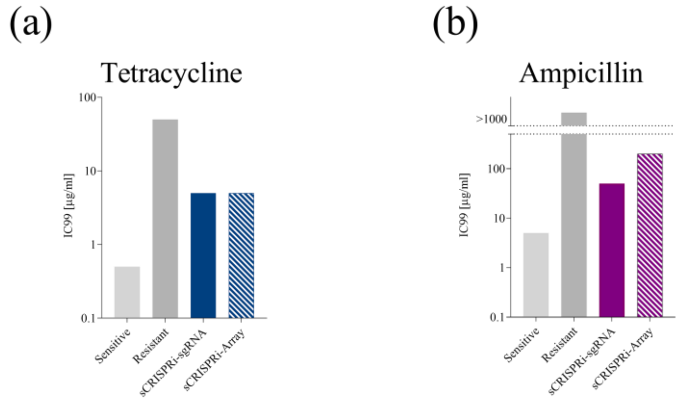

**Figure S7.**

**Agar plate assays for the characterization of CRISPRi-mediated ARG inhibition.**

**(a-b)** Antibiotic concentrations inhibiting >99% bacterial growth ($IC_{99}$) in agar plate assays for tetracycline (a) and ampicillin (b) treated strains. sCRISPRi-sgRNA refers to strains engineered with a two-plasmid CRISPRi platform, including a constitutive dCas9 module and an IPTG-inducible single gRNA module. sCRISPRi-Array refers to strains engineered with a one-plasmid CRISPRi platform, including a HSL-inducible dCas9 module and an IPTG-inducible gRNA module implemented as CRISPRi array. Bars represent $IC_{99}$ recorded for two replicates that provided consistent values, therefore no error bars are present. Data come from experiments in LB media with IPTG or IPTG + HSL=10nM. TOP10, TOP10F', and T-$cr_{sgT\_const}$ and AT-$cr_{arAT}$ were used as *sensitive*, *resistant*, and *sCRISPRi-sgRNA* and *sCRISPRi-Array* strains, respectively, to investigate tetracycline re-sensitization. TOP10F', A-res, and A-$cr_{sgA\_const}$ and AT-$cr_{arAT}$ were used as *sensitive*, *resistant*, and *sCRISPRi-sgRNA* and *sCRISPRi-Array* strains, respectively, to investigate ampicillin re-sensitization.

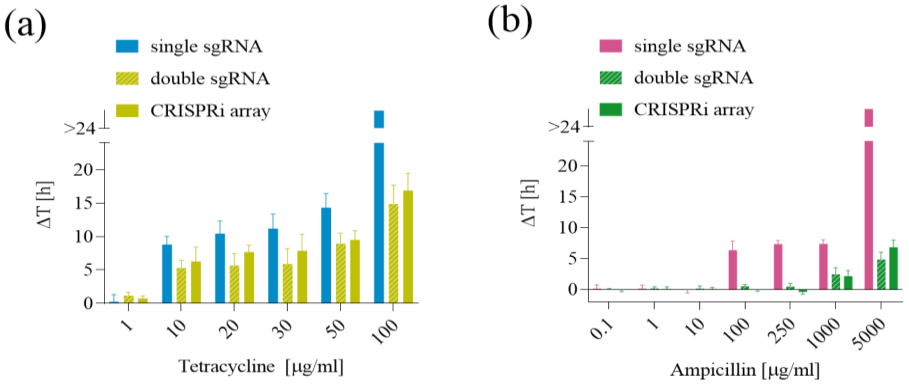

**Figure S8.**

**Comparison of growth delays among single-sgRNA, double-sgRNA, and CRISPRi array designs, with a weak constitutive dCas9 expression cassette in laboratory strains. (a-b)** Growth delays (Δt) of tetracycline (a) and ampicillin (b) *sCRISPRi* strains are shown for the indicated strains relative to the growth profile without antibiotics. *sCRISPRi* strains are engineered with a two-plasmid CRISPRi platform, including constitutive dCas9 and IPTG-inducible single sgRNA cassette, double-sgRNA cassette or CRISPRi array. Bars represent means and standard deviations (N=3). The bars over the interrupted axis indicate the antibiotic concentration (MIC) for which $OD_{600}$ was lower than 0.1. All the data come from experiments in LB media in the presence of IPTG. T-$cr_{sgT\_const}$, A-$cr_{sgA\_const}$ were used as *sCRISPRi* strains with single-sgRNA to investigate tetracycline and ampicillin re-sensitization, respectively. AT-$cr_{sgAT\_const}$ and AT-$cr_{arAT\_const}$ were used as *sCRISPRi* strains with double-sgRNA and CRISPRi array, respectively, to investigate tetracycline and ampicillin re-sensitization.

**Supplementary notes for Figure S9.**

*Multi-targeting of ARGs using an inducible dCas9.* We tuned the intracellular amount of dCas9 by replacing the original constitutive expression module with an N-3-oxohexanoyl-L-homoserine lactone (HSL)-inducible expression cassette, placed in the LC vector, enabling the transition to a one-plasmid platform for ARG repression (Figure 2a, Figure S1a). The resulting *sCRISPRi* strains harbored one LC-plasmid with the full CRISPRi circuit and two different ARG-carrying plasmids. The optimal dCas9 expression level for target gene repression was determined by individual TC and AMP growth inhibition assays in which IPTG was added at a fixed concentration to trigger sgRNA or CRISPRi array transcription, and a range of HSL concentrations were used to tune dCas9 level. No growth defect was detected for increasing HSL concentrations, confirming the non-toxicity of dCas9 expression in the tested range, as previously observed[35]. Among the tested conditions, 10 nM HSL was selected to induce dCas9 in all the experiments below due to its overall superior performance in terms of growth delay, considering TC and AMP experiments with CRISPRi array or tandem sgRNA architectures (Figure 2c,d and Figure S9).

The comparison between CRISPRi array and double sgRNA cassette showed similar growth inhibition performance in terms of MIC with 10 nM HSL (Figure 2d, Figure S9). However, growth delays showed that the CRISPRi array architecture led to a statistically significant increase (20 to 30%) of delay value ($p<0.05$, ANOVA) for sub-lethal antibiotic levels (AMP=5000 µg/ml and TC>10 µg/ml). As expected from the MIC data above, this analysis also showed that delays were significantly higher for the inducible dCas9 condition than the constitutive one for AMP>100 µg/ml ($p<0.05$, ANOVA).

Due to its better performance, we selected the inducible dCas9 with CRISPRi array design for subsequent analyses.

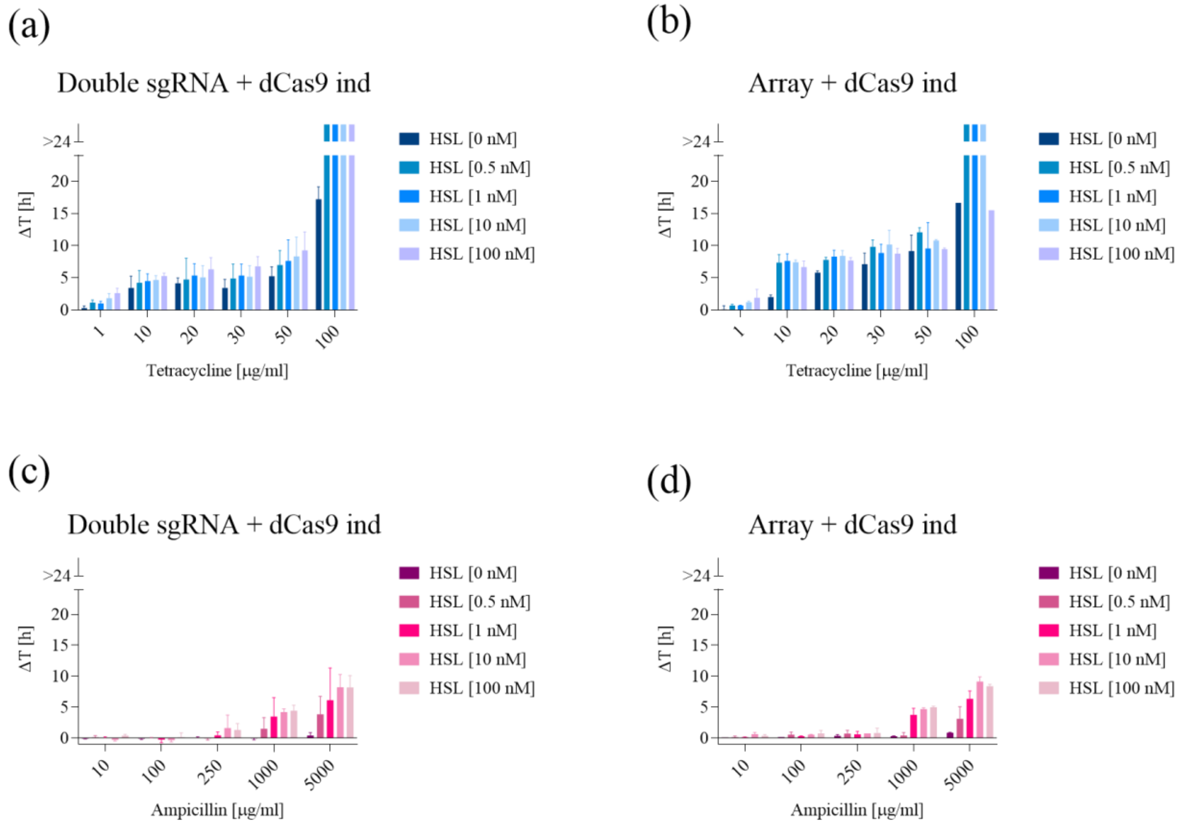

**Figure S9.**

**Growth delays of laboratory strains bearing a one-plasmid CRISPRi platform, including an inducible dCas9 and a two guide RNAs architecture (double sgRNA cassette or CRISPRi array) for different HSL concentrations. (a-d)** Growth delays ($\Delta t$) of tetracycline (a-b) and ampicillin (c-d) *sCRISPRi* strains are shown for the indicated strains relative to the growth profile without antibiotics. *sCRISPRi* strains are engineered with a one-plasmid CRISPRi platform, including HSL-inducible dCas9 and IPTG-inducible double sgRNA cassette (a,c) or CRISPRi array (b,d) transcribing gtetA and $gbla_{promoter}$ simultaneously. Bars represent the mean of at least two independent replicates with error bars indicating standard deviations. The bars over the interrupted axis indicate the antibiotic concentration (MIC) for which cell growth is maintained lower than $OD_{600}$ 0.1 over 24 h. All the data come from experiments in LB media in the presence of IPTG and the indicated HSL concentrations. AT-$cr_{sgAT}$ and AT-$cr_{arAT}$ were used as *sCRISPRi* strains with double-sgRNA and CRISPRi array, respectively, to investigate tetracycline and ampicillin re-sensitization.

### Supplementary notes for Figure S10.

*Inhibition of two ARGs to evaluate a combined antibiotic treatment.* We tested the multi-targeting capability of the designed CRISPRi array by performing microplate assays in which TC and AMP were administered simultaneously to the TC- and AMP-*sCRISPRi* strain. In particular, this test aimed at verifying whether a combined antibiotic treatment could have an advantage over a single antibiotic treatment. This was especially important in case of *bla*$_{\text{TEM-116}}$ repression since the strain individually tested for *bla*$_{\text{TEM-116}}$ inhibition withstood the highest AMP concentration also when engineered with an optimized CRISPRi platform. To counteract this resistance, we evaluated if improvements in growth inhibition could be achieved by providing a second antibiotic, administered at a concentration below its individual MIC. Growth delays for each TC/AMP combination tested are shown in Figure S10: the administration of two antibiotics allowed to increase the recovery of treated cells by 70% compared with the individual treatment with AMP, although regrowth could not be prevented.

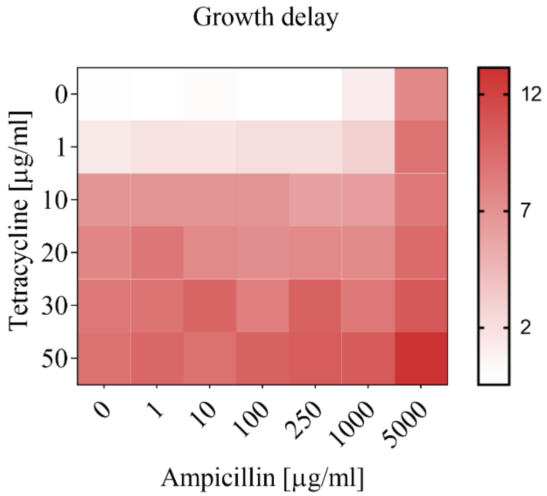

**Figure S10.**

**Growth delays of the laboratory strain bearing a one-plasmid CRISPRi platform including inducible dCas9 and CRISPRi array targeting tetracycline and ampicillin resistance genes, in a combined treatment with the two antibiotics.** Heatmap showing the average growth delay values (Δt) from three independent plate reader experiments with tetracycline and ampicillin at the indicated concentrations, relative to the growth profile without antibiotics. Color scale is shown on the right. All the data come from experiments with the same strain, media and condition as in Figure 2c, d.

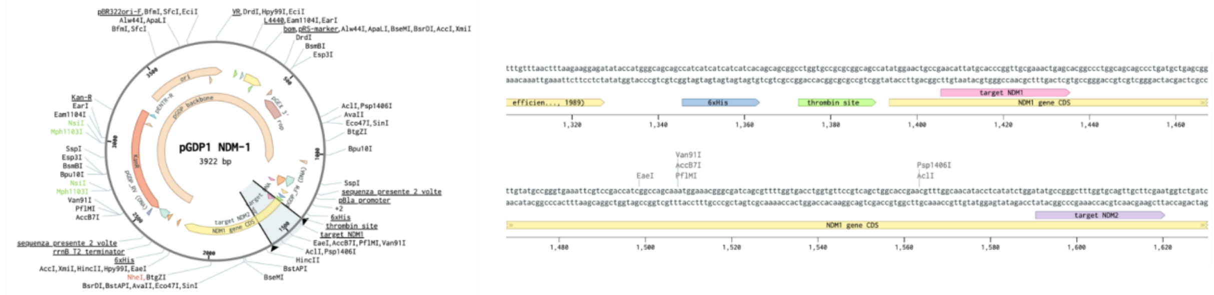

**Figure S11.**

**Target sites of the designed gNDM1 and gNDM2 guides in the *bla*$_{\text{NDM-1}}$ coding sequence.** The codon-optimized *bla*$_{\text{NDM-1}}$ sequence from pGDP1-NDM1 is shown from the Benchling tool. The 20-bp target regions used to design the gNDM1 and gNDM2 spacers are highlighted. The two target sites have a 12- and 197-bp distance from the start codon, respectively.

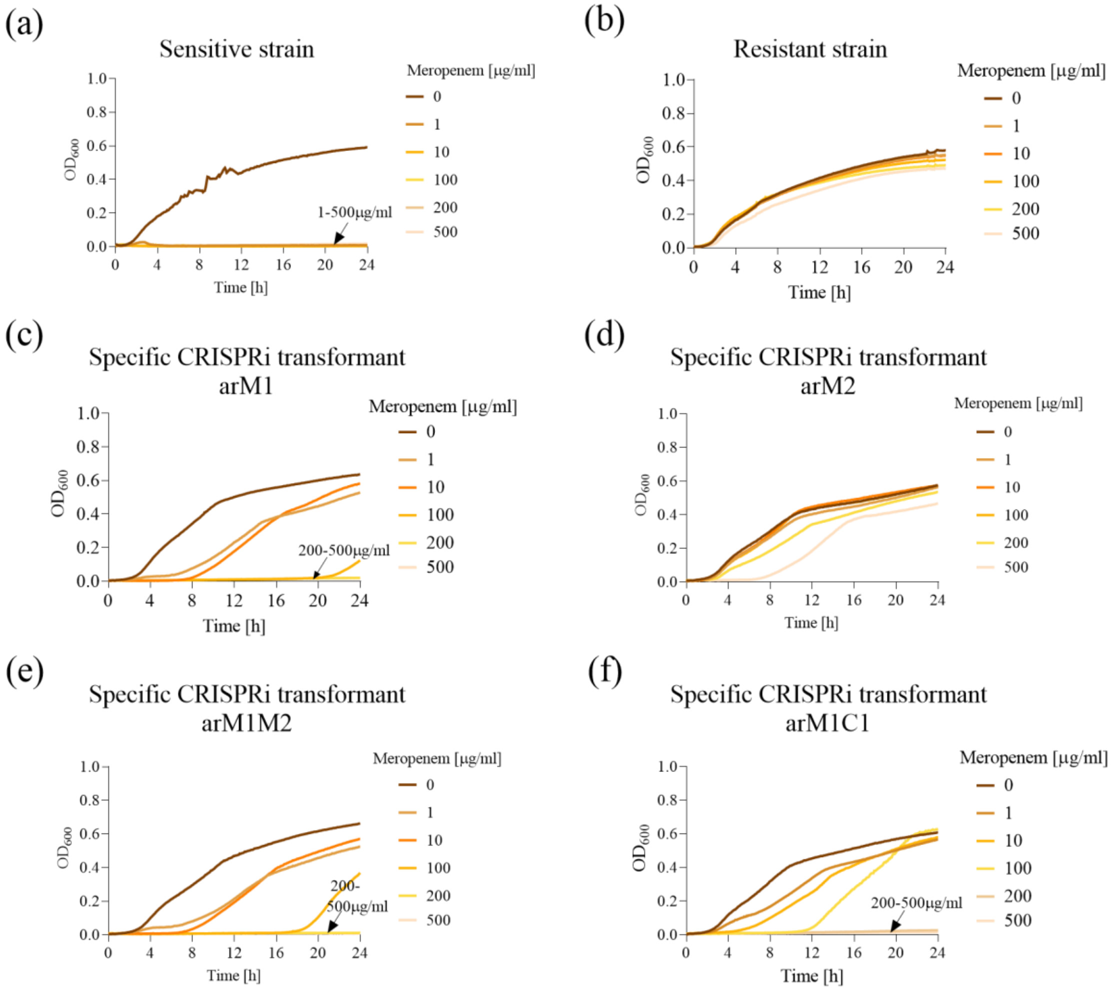

**Figure S12.**

**Re-sensitization to meropenem in laboratory strains using a one-plasmid CRISPRi platform, including HSL-inducible dCas9 and one of four different CRISPRi arrays targeting *bla*$_{\mathbf{NDM\text{-}1}}$. (a-f)** Growth profiles of *sensitive* (a), *resistant* (b) and *sCRISPRi* (c-f) strains from microplate assays to investigate meropenem re-sensitization. *sCRISPRi* strains are engineered with a two-plasmid CRISPRi platform, including HSL-inducible dCas9 and IPTG-inducible CRISPRi array transcribing single gNDM1/gNDM2 (c, d), double gNDM1-gNDM2 (e) and targeting/non-targeting gNDM1-gMCR1 (f) gRNA. Antibiotic concentrations are provided in the legend. The overlapped curves in which a complete growth inhibition was achieved and the list of the corresponding antibiotic concentrations are indicated with an arrow. All the growth profiles are represented as optical density at 600 nm ($OD_{600}$) over time. Representative growth curves are shown from a set of at least three independent experiments. All the data come from experiments in LB media in the presence of IPTG and HSL. TOP10F', M-res and M-$cr_{arM1}$/M-$cr_{arM2}$/ M-$cr_{arM1M2}$/M-$cr_{arM1C1}$ were used as *sensitive*, *resistant* and *sCRISPRi* strains, respectively, to investigate meropenem re-sensitization.

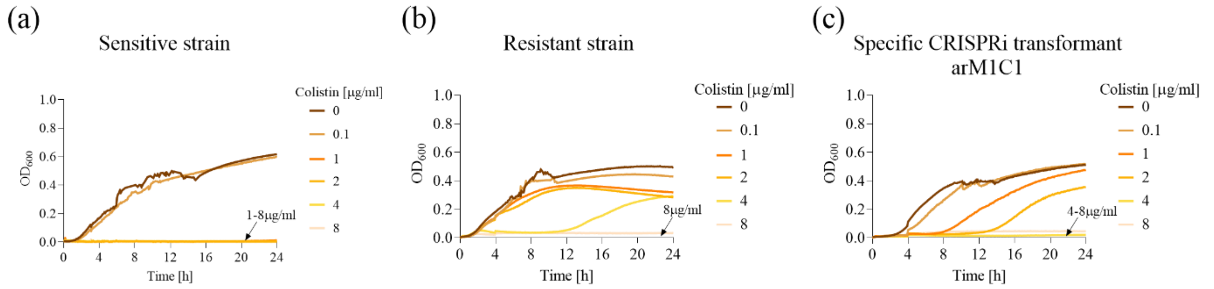

**Figure S13.**

**Re-sensitization to colistin in laboratory strains using a one-plasmid CRISPRi platform, including HSL-inducible dCas9 and CRISPRi array targeting *mcr-1*. (a-c)** Growth profiles of *sensitive* (a), *resistant* (b) and *sCRISPRi* (c) strains from microplate assays to investigate colistin re-sensitization. *sCRISPRi* strain is engineered with a two-plasmid CRISPRi platform, including HSL-inducible dCas9 and IPTG-inducible CRISPRi array transcribing gMCR1. Antibiotic concentrations are provided in the legend. The overlapped curves in which a complete growth inhibition was achieved and the list of the corresponding antibiotic concentrations are indicated with an arrow. All the growth profiles are represented as optical density at 600 nm ($OD_{600}$) over time. All the data come from experiments in LB media. TOP10F', C-res and C-cr$_{arM1C1}$ were used as *sensitive*, *resistant* and *sCRISPRi* strains, respectively, to investigate colistin re-sensitization.

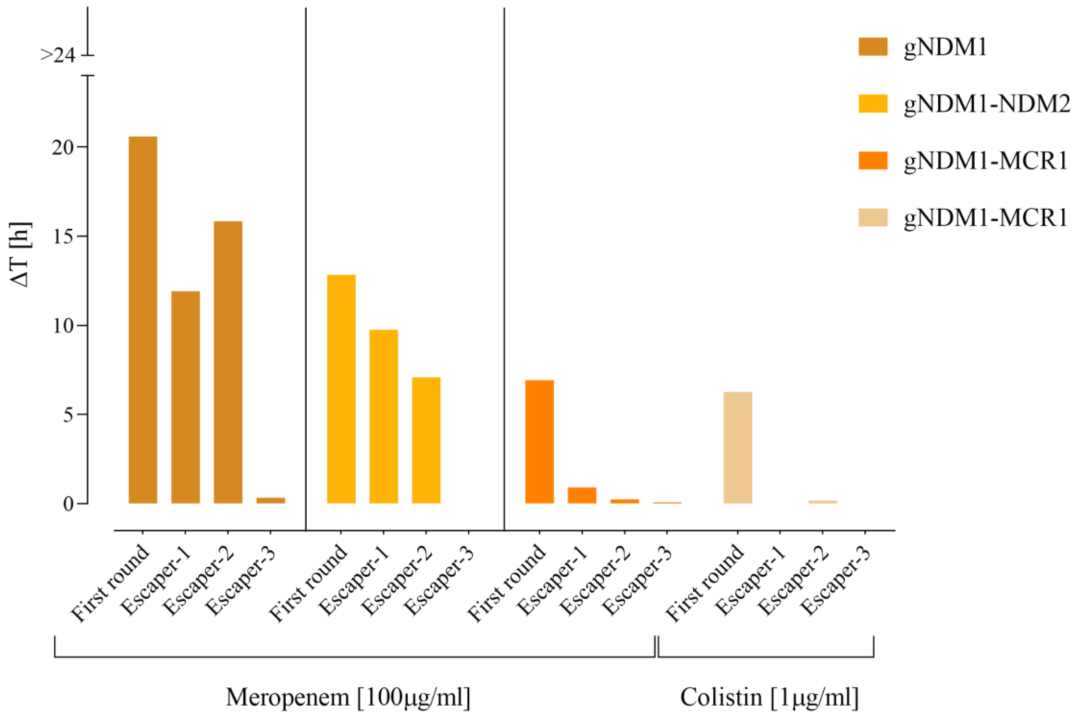

**Figure S14.**

**Growth delays of escaper mutants during second-round experiments**. The order of strains is consistent with Figure 3 in the main text. The delays of the first-round experiments are shown for comparison. All the data come from experiments in LB media in the presence of IPTG and HSL.

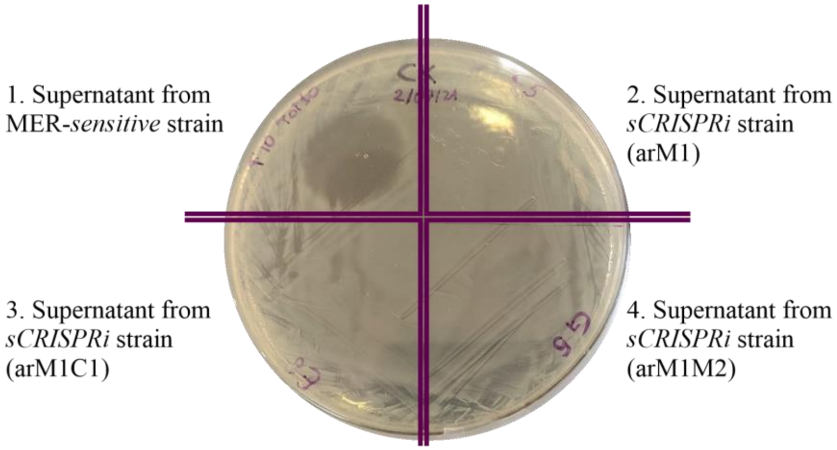

**Figure S15.**

**Supernatant assay with *bla*$_{\mathrm{NDM\text{-}1}}$-expressing laboratory strains.** Supernatants (3 µl) of 24 h lab strain cultures from a meropenem microplate assay were spotted on an LB agar plate in which MER-*sensitive* bacteria were plated. The supernatants were from four strains assayed in presence of 100 µg/ml MER: a *sensitive* strain and three *sCRISPRi* strains (M-cr$_{\mathrm{arM1}}$, M-cr$_{\mathrm{arM1M2}}$ and M-cr$_{\mathrm{arM1C1}}$) in which the growth of escapers was observed.

### Supplementary notes for Figure S16

*Design of bacterial conjugation platform*. To address the delivery of CRISPRi into target cells, we selected bacterial conjugation due to its potential application to *in vivo* therapies, and also to *in vitro* research studies, to enable the incorporation of CRISPRi-plasmids in hard-to-transform strains like clinical isolates[48]. In particular, we engineered *E. coli* donor cells with a *trans*-conjugative platform based on the IncP2 plasmid[49]. The platform consists of two plasmids: the pTA-Mob helper plasmid, bearing the conjugative machinery, and a mobilizable vector carrying the CRISPRi circuitry. The mobilizable vector was constructed by assembling an origin of transfer (oriT from pRK2) into the chloramphenicol-resistant LC plasmids with HSL-inducible dCas9 and IPTG-inducible CRISPRi array targeting single ARGs (Figure S16a). Recognition of the oriT sequence by the relaxase expressed *in trans* from pTA-Mob allowed the CRISPRi-vector to be transferred into antibiotic-resistant recipient cells, which become stable transconjugants.

*Characterization of bacterial conjugation platform.* Conjugations were performed using donor cells engineered with a CRISPRi array targeting *bla*$_{\mathrm{TEM\text{-}116}}$, *bla*$_{\mathrm{NDM\text{-}1}}$, or *mcr-1* gene (strain Donor$_{\mathrm{arAT}}$, Donor$_{\mathrm{arM1}}$, Donor$_{\mathrm{arM1C1}}$). Conjugation frequency was determined after a 24 h mating. We also quantified the ability of the transferred plasmid to increase antibiotic susceptibility in the generated transconjugants. For conjugations with the A-res recipient strain, expressing no auxiliary antibiotic resistance in the ARG-bearing plasmid, we measured the concentration of transconjugants (CHL + AMP) and survivors (CHL + AMP + IPTG/HSL) to ampicillin administration when the CRISPRi circuitry was induced. For conjugations with M- and C-res recipient strains, we took advantage from the presence of a kanamycin resistance cassette (auxiliary antibiotic resistance in the same ARG-bearing plasmid) to discriminate between transconjugants (CHL + KAN) and survivors (CHL + MER/COL + IPTG/HSL) to meropenem or colistin administration in presence of the same inducers.

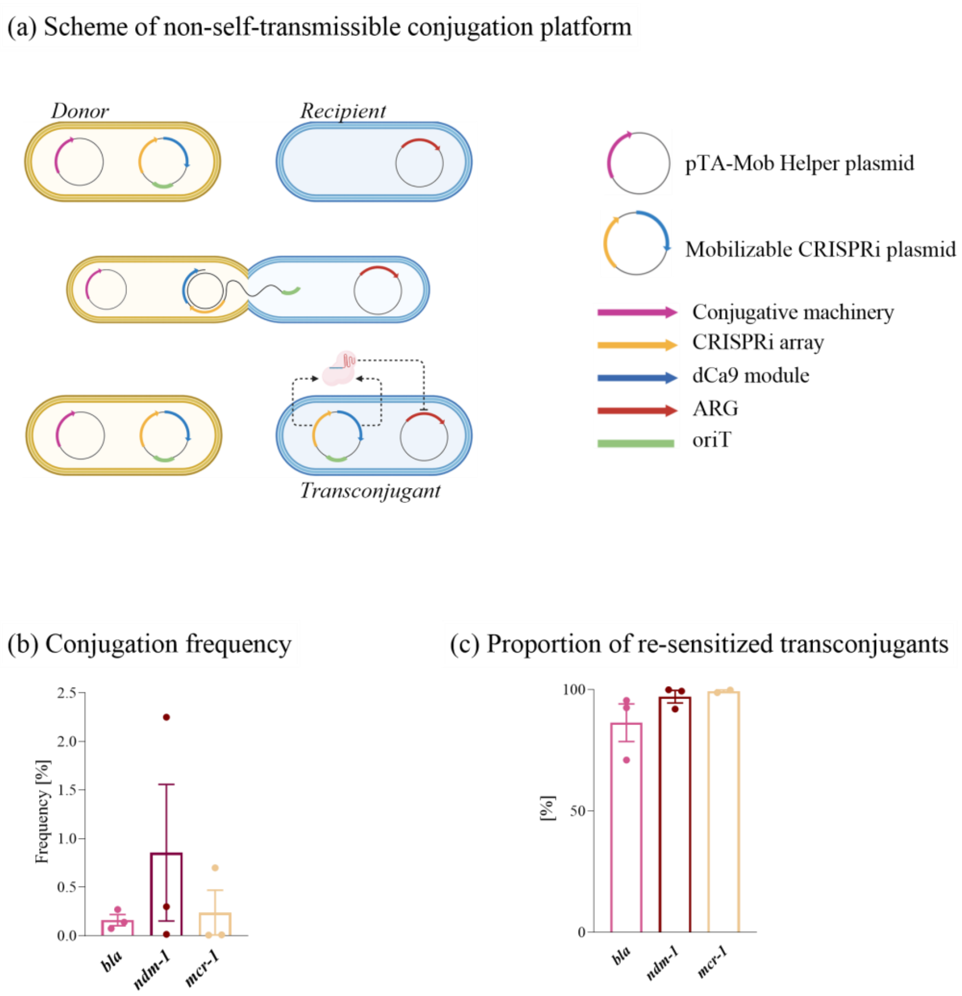

**Figure S16.**

**Horizontal transfer of the CRISPRi plasmid from a donor strain to antibiotic-resistant laboratory strains. (a)** Illustration of conjugative transfer, showing the involved plasmids and the resulting plasmid configuration in transconjugant strains. **(b)** Conjugation efficiency in the three indicated antibiotic-resistant strains. **(c)** Proportion of re-sensitized transconjugant population in the three indicated antibiotic-resistant strains. In panels (b-c), bars indicate the mean of the represented data points (circles), and error bars represent their standard deviations.

**Supplementary notes for Figure S17**

MH plate assays were performed with *sCRISPRi*$^{trans}$ and *nsCRISPRi*$^{trans}$ for investigating meropenem and colistin re-sensitization in two *E. coli* clinical isolates bearing *bla*$_{NDM-5}$ and *mcr-1* gene.

We measured the concentration of transconjugants (CHL + LEV/CIP, see Methods section for details on transconjugant selection) and survivors (CHL + MER/COL + IPTG/HSL) to meropenem or colistin administration. Results confirmed the impact of CRISPRi silencing on the expression of both target genes since the concentration of surviving transconjugants decreased up to 163- and 4-fold for MER 1 µg/ml and COL 0.25 µg/ml, respectively (Figure S17). For both antibiotics, in treated strains the colony counts were significantly lower for specific CRISPRi than non-specific controls ($p<0.05$, t-test). Conversely, strains without antibiotics had a comparable number of viable cells varying less than 1.2-fold with no statistical difference (Figure S17).

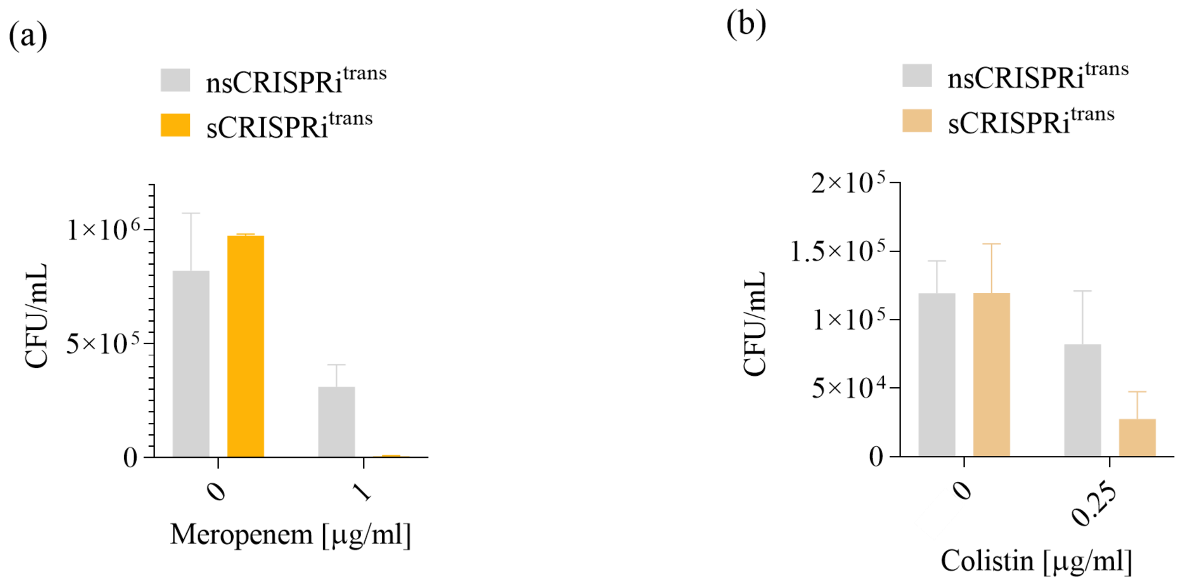

**Figure S17.**

**Meropenem and colistin re-sensitization in agar plate assays for two clinical isolates with *bla*$_{\mathbf{NDM\text{-}5}}$ and *mcr-1*. (a-b)** Colony forming units (CFUs) are shown for the non-specific (*nsCRISPRi*$^{trans}$) and specific (*sCRISPRi*$^{trans}$) CRISPRi strains bearing *bla*$_{NDM\text{-}5}$ (a) or *mcr-1* (b) target gene, after conjugations in which the transconjugants were selected in absence and presence of the indicated antibiotic. Bars represent the average value of two (*ndm-5* strains) or three (*mcr-1* strains) independent experiments, with error bars representing standard deviations. M-cr$^{CI}_{arAT}$ and M-cr$^{CI}_{arM(w)}$ were used as *nsCRISPRi*$^{trans}$ and *sCRISPRi*$^{trans}$, respectively, to investigate meropenem re-sensitization. C-cr$^{CI}_{arAT}$ and C-cr$^{CI}_{arM1C1}$ were used as *nsCRISPRi*$^{trans}$ and *sCRISPRi*$^{trans}$, respectively, to investigate colistin re-sensitization.

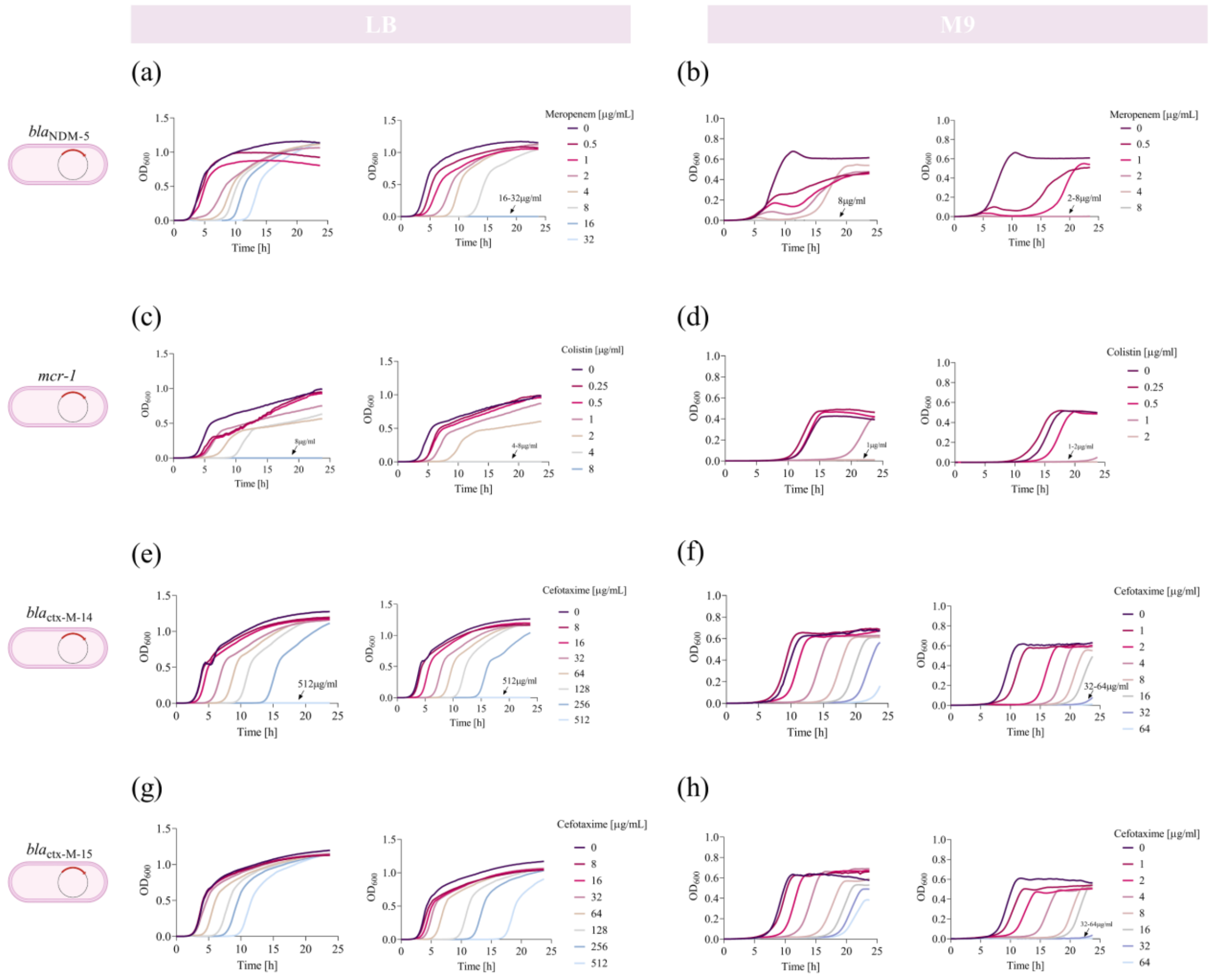

**Figure S18.**

**Antibiotic re-sensitization of four *E. coli* clinical isolates grown in LB and M9 media. (a-h)** Growth profiles and delays of *nsCRISPRi*$^{trans}$ (subpanel on the left) and *sCRISPRi*$^{trans}$ (subpanels on the right) from microplate assays to investigate meropenem (a-b), colistin (c-d) and cefotaxime (e-h) re-sensitization. *sCRISPRi*$^{trans}$ are engineered with the one-plasmid CRISPRi platform described in Figure 4. Data in panels (a,c,e,g) were obtained from LB medium microplate assays, and data in panels (b,d,f,h) were obtained from minimal M9 microplate assays. Representative growth curves are shown from a set of at least three independent experiments. Data come from experiments in LB medium in the presence of IPTG and HSL, or M9 medium with lactose in the presence of HSL. Strains used to investigate meropenem, colistin and cefotaxime re-sensitization are listed in the caption of Figure 4.

### Supplementary Tables

#### Table S1.

List of strains used in this work. LC, MC and HC (pSB4C5, pSB3K3 and pSB1A2) vector backbones carrying a collection of synthetic constructs (also listed in Table S2) were inserted in TOP10, TOP10F' and DH10B host strains. Endogenous plasmids were already present (*E. coli* clinical isolates) or previously incorporated (pTA-Mob, F') in the host strain. The CRISPRi targeted gene is reported, along with a description of the designed function for each strain used in this work. In the "description" section, antibiotic resistances used as auxiliary markers for transconjugant selection are underlined, while those used as CRISPRi targets are written in bold type. NA=not available.

| **Strain** | **Host** | **LC** | **MC** | **HC** | **Endogenous plasmids** | **Target** | **Description** |
|---|---|---|---|---|---|---|---|
| TOP10 | TOP10 | - | - | - | - | - | TC- susceptible control |
| TOP10 F' | TOP10 F' | - | - | - | F' episome | - | TC- resistant control; AMP-, MER-, COL- susceptible control |
| T-cr$_{sgA_const}$ | TOP10 F' | gbla$_{CDS}$ | J116dCas9 | - | F' episome | - | TC-non-specific CRISPRi transformant with non-targeting gRNA |
| T-cr$_{sgT_const}$ | TOP10 F' | gtetA | J116dCas9 | - | F' episome | *tetA* | TC-specific CRISPRi transformant with gtetA targeting tetA CDS |
| A-res | TOP10 F' | - | - | I13521 | F' episome | - | AMP-resistant control and recipient for conjugation with Donor$_{arAT}$; **AMP$^{R}$** |
| A-cr$_{sgT_const}$ | TOP10 F' | gtetA | J116dCas9 | I13521 | F' episome | - | AMP-non-specific CRISPRi transformant with non-targeting gRNA |
| A-cr$_{sgA_const}$ | TOP10 F' | gbla$_{promoter}$ | J116dCas9 | I13521 | F' episome | *bla*$_{TEM-116}$ | AMP-specific CRISPRi transformant with gbla$_{promoter}$ targeting P$_{bla}$ promoter |
| A-cr$_{sgA(CDS)_const}$ | TOP10 F' | gbla$_{CDS}$ | J116dCas9 | I13521 | F' episome | *bla*$_{TEM-116}$ | AMP-specific CRISPRi transformant with gbla$_{CDS}$ targeting *bla*$_{TEM-116}$ CDS |
| AT-cr$_{sgAT_const}$ | TOP10 F' | gbla-gtetA$_{cassette}$ | J116dCas9 | I13521 | F' episome | *bla*$_{TEM-116}$/*tetA* | AMP- and TC- specific CRISPRi transformant with gbla$_{promoter}$ targeting P$_{bla}$ promoter and gtetA targeting *tetA* gene |

| | | | | | | | |
|---|---|---|---|---|---|---|---|
| AT-$cr_{arAT\_const}$ | TOP10 F' | gbla-$gtetA_{array}$ | J116dCas9 | I13521 | F' episome | *bla*$_{TEM-116}$/*tetA* | AMP- and TC- specific CRISPRi transformant with $gbla_{promoter}$ targeting $P_{bla}$ promoter and gtetA targeting *tetA* gene |
| AT-$cr_{sgAT}$ | TOP10 F' | gbla-$gtetA_{cassette}$ + dCas9 | - | I13521 | F' episome | *bla*$_{TEM-116}$/*tetA* | AMP- and TC- specific CRISPRi transformant with $gbla_{promoter}$ targeting $P_{bla}$ promoter and gtetA targeting *tetA* gene |
| AT-$cr_{arAT}$ | TOP10 F' | gbla-$gtetA_{array}$ + dCas9 | - | I13521 | F' episome | *bla*$_{TEM-116}$/*tetA* | AMP- and TC- specific CRISPRi transformant with $gbla_{promoter}$ targeting $P_{bla}$ promoter and gtetA targeting *tetA* gene |
| M-res | TOP10 F' | - | pGDP1 NDM-1 | - | F' episome | - | MER-resistant control and recipient for conjugation with $Donor_{arM1}$ ; $KAN^{R}$, **$MER^{R}$** |
| M-$cr_{arM1}$ | TOP10 F' | oriT_$gNDM1_{array}$ + dCas9 | pGDP1 NDM-1 | - | F' episome | *bla*$_{NDM-1}$ | MER-specific CRISPRi transformant with gNDM1 targeting *bla*$_{NDM-1}$ CDS at site 1 |
| M-$cr_{arM2}$ | TOP10 F' | oriT_$gNDM2_{array}$ + dCas9 | pGDP1 NDM-1 | - | F' episome | *bla*$_{NDM-1}$ | MER-specific CRISPRi transformant with gNDM2 targeting *bla*$_{NDM-1}$ CDS at site 2 |
| M-$cr_{arM1M2}$ | TOP10 F' | oriT_gNDM1-$gNDM2_{array}$ + dCas9 | pGDP1 NDM-1 | - | F' episome | *bla*$_{NDM-1}$ | MER-specific CRISPRi transformant with gNDM1 and gNDM2 targeting *bla*$_{NDM-1}$ CDS at site 1 and site 2 |
| M-$cr_{arM1C1}$ | TOP10 F' | oriT_gNDM1-$gMCR1_{array}$ + dCas9 | pGDP1 NDM-1 | - | F' episome | *bla*$_{NDM-1}$ | MER-specific CRISPRi transformant with gNDM1 targeting *bla*$_{NDM-1}$ CDS at site 1 |
| C-res | TOP10 F' | - | pGDP2 MCR-1 | - | F' episome | - | COL-resistant control and recipient for conjugation with $Donor_{arM1C1;}$ $KAN^{R}$, **$COL^{R}$** |

| | | | | | | | |
|---|---|---|---|---|---|---|---|
| C-cr$_{arM1C1}$ | TOP10 F' | gNDM1-gMCR1$_{array}$ + dCas9 | pGDP2 MCR-1 | - | F' episome | *mcr-1* | COL-specific CRISPRi transformant with gMCR1 targeting *mcr-1* CDS |
| Donor$_{arAT}$ | DH10B | oriT_gAMP-gtetA$_{array}$ + dCas9 | - | - | p TA-Mob | - | Donor for conjugation with A-res, M-res$^{CI}$, C-res$^{CI}$ |
| Donor$_{arM1}$ | DH10B | oriT_gNDM1$_{array}$ + dCas9 | - | - | p TA-Mob | - | Donor for conjugation with M-res |
| Donor$_{arM1C1}$ | DH10B | oriT_gNDM1-gMCR1$_{array}$ + dCas9 | - | - | p TA-Mob | - | Donor for conjugation with C-res and C-res$^{CI}$ |
| A-cr$_{arAT}$ $^{trans}$ | TOP10 F' | oriT_gbla-gtetA$_{array}$ + dCas9 | - | I13521 | - | *bla*$_{TEM-116}$ | AMP-specific CRISPRi transconjugant with with gbla$_{promoter}$ targeting P$_{bla}$ promoter |
| M-cr$_{arM1}$$^{trans}$ | TOP10 F' | oriT_gNDM1$_{array}$ + dCas9 | pGDP1 NDM-1 | - | - | *bla*$_{NDM-1}$ | MER-specific CRISPRi transconjugant with gNDM1 targeting *bla*$_{NDM-1}$ CDS at site 1 |
| C-cr$_{arM1C1}$$^{trans}$ | TOP10 F' | oriT_gNDM1-gMCR$_{array}$ + dCas9 | pGDP2 MCR-1 | - | - | *mcr-1* | COL-specific CRISPRi transconjugant with gMCR1 targeting *mcr-1* CDS |
| Donor$_{arM(wt)}$ | DH10B | oriT_gNDM(wt) + dCas9 | - | - | p TA-Mob | - | Donor for conjugation with M-res$^{CI}$, X14-res$^{CI}$, X15-res$^{CI}$ |
| Donor$_{arX14}$ | DH10B | oriT_gCTX-M-14 + dCas9 | - | - | p TA-Mob | - | Donor for conjugation with X14-res$^{CI}$ |
| Donor$_{arX15}$ | DH10B | oriT_gCTX-M-15 + dCas9 | - | - | p TA-Mob | - | Donor for conjugation with X15-res$^{CI}$ |

| | | | | | | | |
|---|---|---|---|---|---|---|---|
| M-res$^{CI}$ | - | - | - | - | *bla*$_{NDM-5}$ bearing plasmid | - | MER-resistant recipient for conjugation with Donor$_{arM(wt);}$ LEV$^R$, **MER$^R$** |
| M-cr$^{CI}_{arAT}$ | - | oriT_gbla-gtetA$_{array}$ + dCas9 | - | - | *bla*$_{NDM-5}$ bearing plasmid | - | MER (wt)-non-specific CRISPRi transconjugant with non-targeting gRNA |
| M-cr$^{CI}$ $_{arM(wt)}$ | - | oriT_gNDM(wt) + dCas9 | - | - | *bla*$_{NDM-5}$ bearing plasmid | *bla*$_{NDM-5}$ | MER (wt)-specific CRISPRi transconjugant with gNDM(wt) targeting *bla*$_{NDM-5}$ CDS |
| C-res$^{CI}$ | - | - | - | - | NA | - | Recipient for conjugation with Donor$_{arM1C1}$, CIP$^R$, **COL$^R$** |
| C-cr$^{CI}$ $_{arAT}$ | - | oriT_gbla-gtetA$_{array}$ + dCas9 | - | - | NA | - | COL (wt)-non-specific CRISPRi transconjugant with non-targeting gRNA |
| C-cr$^{CI}$ $_{arM1C1}$ | - | oriT_gNDM1-gMCR1$_{array}$ + dCas9 | - | - | NA | *mcr-1* | COL (wt)-specific CRISPRi transconjugant with gMCR1 targeting *mcr-1* CDS |
| X14-res$^{CI}$ | - | - | - | - | Chromosome | - | Recipient for conjugation with Donor$_{arX14}$ |
| X14-cr$^{CI}$ $_{arM(wt)}$ | - | oriT_gNDM1$_{array}$ + dCas9 | - | - | Chromosome | - | CTX-M-14-non-specific CRISPRi transconjugant with non-targeting gRNA |
| X14-cr$^{CI}$ $_{arX14}$ | - | oriT_gCTX-M-14 + dCas9 | - | - | Chromosome | *bla*$_{ctx-M-14}$ | CTX-M-14-specific CRISPRi transconjugant with gCTX-M-14 targeting *bla*$_{ctx-M-14}$ CDS |

| | | | | | | | |
|---|---|---|---|---|---|---|---|
| X15-res$^{CI}$ | - | - | - | - | Chromosome | - | Recipient for conjugation with Donor$_{arX15}$ |
| X15-cr$^{CI}$ $_{arM(wt)}$ | - | oriT_gNDM1$_{array}$ + dCas9 | - | - | Chromosome | - | CTX-M-15-non-specific CRISPRi transconjugant with non-targeting gRNA |
| X15-cr$^{CI}$ $_{arX15}$ | - | oriT_gCTX-M-15 + dCas9 | - | - | Chromosome | *bla*$_{ctx\text{-}M\text{-}15}$ | CTX-M-15-specific CRISPRi transconjugant with gCTX-M-15 targeting *bla*$_{ctx\text{-}M\text{-}15}$ CDS |

**Table S2.**
List of recombinant plasmids used in this work. The replication origin, the plasmid code and a description of the genetic parts composing each plasmid are reported. Plasmid codes are relative to the MIT Registry of Standard Biological Parts (http://parts.igem.org), in which sequences are available; the BBa_ prefix is omitted. These codes indicate the inserts of the LC, MC and HC (pSB4C5, pSB3K3 and pSB1A2) vector backbones.

| **Plasmid** | **Copy Number** | **Replication origin** | **Code** | **Description** |
|---|---|---|---|---|
| I3521 | HC | mutated pMB1 | pSB1A2-I13521 | Recombinant vector carrying the $bla_{\text{TEM-116}}$ gene under the $P_{\text{bla}}$ promoter |
| J116dCas9 | MC | p15A | pSB3K3-J107202 | Recombinant vector carrying a constitutive dCas9 cassette under the $P_{\text{J23116}}$ promoter |
| pGDP1-NDM1 | MC | pBR322 | Addgene n.#112883 | Recombinant vector carrying a codon-optimized version of the $bla_{\text{NDM-1}}$ gene under the $P_{\text{bla}}$ promoter |
| pGDP2-MCR1 | MC | pBR322 | Addgene n.#118404 | Recombinant vector carrying *mcr-1* gene under the $P_{\text{Llac01}}$ promoter |
| gtetA | LC | pSC101 | pSB4C5-J107161 | IPTG-inducible single sgRNA cassette transcribing gtetA under the $P_{\text{Llac01}}$ promoter |
| $\text{gbla}_{\text{promoter}}$ | LC | pSC101 | pSB4C5-J107162 | IPTG-inducible single sgRNA cassette transcribing $\text{gbla}_{\text{promoter}}$ under the $P_{\text{Llac01}}$ promoter |
| $\text{gbla}_{\text{CDS}}$ | LC | pSC101 | pSB4C5-J107163 | IPTG-inducible single sgRNA cassette transcribing $\text{gbla}_{\text{CDS}}$ under the $P_{\text{Llac01}}$ promoter |
| $\text{gbla-gtetA}_{\text{cassette}}$ | LC | pSC101 | pSB4C5-J107164 | IPTG-inducible double sgRNA cassette transcribing $\text{gbla}_{\text{promoter}}$ and gtetA under the $P_{\text{Llac01}}$ promoter |
| $\text{gbla-gtetA}_{\text{array}}$ | LC | pSC101 | pSB4C5-J107165 | IPTG-inducible CRISPRi array transcribing $\text{gbla}_{\text{promoter}}$ and gtetA under the $P_{\text{Llac01}}$ promoter |

| | | | | |
|---|---|---|---|---|
| gbla-gtetA$_{cassette}$ + dCas9 | LC | pSC101 | pSB4C5-J107166 | IPTG-inducible double sgRNA cassette transcribing gbla$_{promoter}$ and gtetA under the $P_{Llac01}$ promoter + HSL-inducible dCas9 cassette under $P_{lux}$ promoter |
| gbla-gtetA$_{array}$ + dCas9 | LC | pSC101 | pSB4C5-J107167 | IPTG-inducible CRISPRi array transcribing gbla$_{promoter}$ and gtetA under the $P_{Llac01}$ promoter + HSL-inducible dCas9 cassette under the $P_{lux}$ promoter |
| oriT_gNDM1$_{array}$ + dCas9 | LC | pSC101 | pSB4C5-J107168 | IPTG-inducible CRISPRi array transcribing gNDM1 under the $P_{Llac01}$ promoter + HSL-inducible dCas9 under the $P_{lux}$ promoter, with the origin of transfer sequence (oriT) from pRK2 |
| oriT_gNDM2$_{array}$ + dCas9 | LC | pSC101 | pSB4C5-J107169 | IPTG-inducible CRISPRi array transcribing gNDM2 under the $P_{Llac01}$ promoter + HSL-inducible dCas9 under the $P_{lux}$ promoter, with the origin of transfer sequence (oriT) from pRK2 |
| oriT_gNDM1-gNDM2$_{array}$ + dCas9 | LC | pSC101 | pSB4C5-J107170 | IPTG-inducible CRISPRi array transcribing gNDM1 and gNDM2 under the $P_{Llac01}$ promoter + HSL-inducible dCas9 under the $P_{lux}$ promoter, with the origin of transfer sequence (oriT) from pRK2 |
| oriT_gNDM1-gMCR1$_{array}$ + dCas9 | LC | pSC101 | pSB4C5-J107171 | IPTG-inducible CRISPRi array transcribing gNDM1 and gMCR1 under the $P_{Llac01}$ promoter + HSL-inducible dCas9 under the $P_{lux}$ promoter, with the origin of transfer sequence (oriT) from pRK2. |
| oriT_gbla-gtetA$_{array}$ + dCas9 | LC | pSC101 | pSB4C5-J107172 | Same as gbla-gtetA$_{array}$ + dCas9, with the origin of transfer sequence (oriT) from pRK2 |
| oriT_gNDM(wt) + dCas9 | LC | pSC101 | pSB4C5-J107175 | IPTG-inducible CRISPRi array transcribing gNDM$_{(wt)}$ under the $P_{Llac01}$ promoter + HSL-inducible dCas9 under the $P_{lux}$ promoter. gNDM(wt) targets the wt sequence of the *bla*$_{NDM-5}$ gene |
| oriT_gCTX-M-14 + dCas9 | LC | pSC101 | pSB4C5-J107176 | IPTG-inducible CRISPRi array transcribing gCTX-M-14 under the $P_{Llac01}$ promoter + HSL-inducible dCas9 under the $P_{lux}$ promoter. |
| oriT_gCTX-M-15 + dCas9 | LC | pSC101 | pSB4C5-J107177 | IPTG-inducible CRISPRi array transcribing gCTX-M-15 under the $P_{Llac01}$ promoter + HSL-inducible dCas9 under the $P_{lux}$ promoter. |

**Table S3.**
List of primers used in this work for escaper analysis. For each template, the primer pairs used for DNA sequencing are reported along with a description of the specific part to be sequenced.

| Template | Primer Pair | Sequence (5'-3') | Description |
|---|---|---|---|
| gNDM1 / gNDM1-gNDM2 / gNDM1-gMCR1$_{array}$ + dCas9 | C0062_VF | gaatgtttagcgtgggcatg | Sequencing of CRISPRi array and dCas9 gene from the circuits encoding *bla*$_{NDM-1}$ and *mcr-1* spacers |
| | FW_dCas9 | caatccatcactggtctttatg | |
| pGDP1 NDM-1 | FW_pGDP | cgtatcggtgattcattctgc | Sequencing of *bla*$_{NDM-1}$ and *mcr-1* gene |
| pGDP2 MCR-1 | RV_pGDP | gcagtttcatttgatgctcgat | |

**Table S4.**
List of IC$_{99}$ values determined with agar plate assays. All the values are expressed as (µg/mL).

| | **Sensitive** | **Resistant** | **Specific CRISPRi transformant** | | | | |
|---|---|---|---|---|---|---|---|
| **Antibiotic [µg/mL]** | | | single sgRNA | sgRNA cassette + dCas9 const | Array + dCas9 const | sgRNA cassette + dCas9 ind | Array + dCas9 ind |
| **TC** | 0.5 | 50 | 5 | / | / | / | 5 |
| **AMP** | 5 | > 1000 | 50 (gbla$_{promoterr}$)<br>150 (gbla$_{CDS}$) | / | / | / | 200 |
| **MER** | 0.0312 | 10 | / | / | / | / | 0.125 |
| **COL** | 0.0312 | 2 | / | / | / | / | 0.125 |
